# Space-time dynamics in monitoring neotropical fish communities using eDNA metabarcoding

**DOI:** 10.1101/2020.02.04.933366

**Authors:** Naiara Guimarães Sales, Owen Simon Wangensteen, Daniel Cardoso Carvalho, Kristy Deiner, Kim Præbel, Ilaria Coscia, Allan D. McDevitt, Stefano Mariani

**Affiliations:** Ecosystems and Environment Research Centre, School of Science, Engineering and Environment, University of Salford, UK; CESAM – Centre for Environmental and Marine Studies, Departamento de Biologia Animal, Faculdade de Ciências da Universidade de Lisboa, Lisbon, Portugal; Norwegian College of Fishery Science, UiT - The Arctic University of Norway, Tromsø, Norway; Programa de Pós-graduação em Biologia de Vertebrados, Pontifícia Universidade Católica de Minas Gerais, Belo Horizonte, Brasil; Life Sciences, Natural History Museum, London, UK; School of Natural Sciences and Psychology, Liverpool John Moores University, Liverpool, UK

**Keywords:** eDNA, biodiversity assessment, fish, freshwater, Brazil, river

## Abstract

The biodiverse Neotropical ecoregion remains insufficiently assessed, poorly managed, and threatened by unregulated human activities. Novel, rapid and cost-effective DNA-based approaches are valuable to improve understanding of the biological communities and for biomonitoring in remote areas. Here, we evaluate the potential of environmental DNA (eDNA) metabarcoding for assessing the structure and distribution of fish communities by analysing sediments and water from 11 locations along the Jequitinhonha River catchment (Brazil). Each site was sampled twice, before and after a major rain event in a five-week period and fish diversity was estimated using high-through-put sequencing of 12S rRNA amplicons. In total, 252 Molecular Operational Taxonomic Units (MOTUs) and 34 fish species were recovered, including endemic, introduced, and previously unrecorded species for this basin. Spatio-temporal variation of fish assemblages was detected, richness during the first campaign was nearly twice as high as in the second sampling round; though peaks of diversity were primarily associated with only four locations. No correlation between β-diversity and longitudinal distance or presence of dams was detected, but low species richness observed at sites located near dams indicates that these anthropogenic barriers might have an impact on local fish diversity. Unexpectedly high α-diversity levels recorded at the river mouth suggest that these sections should be further evaluated as putative “eDNA reservoirs” for rapid monitoring. By uncovering spatio-temporal changes, unrecorded biodiversity components, and putative anthropogenic impacts on fish assemblages, we further strengthen the potential of eDNA metabarcoding as a biomonitoring tool, especially in regions often neglected or difficult to access.

## 1 INTRODUCTION

Despite covering less than 1% of the Earth’s surface, freshwater habitats harbour over 40% of global fish diversity (Nelson, 2006; Dudgeon et al., 2006). Fish from rivers, lakes, and wetlands provide essential protein subsistence for a large proportion of human populations worldwide (FAO, 2012; McIntyre et., 2016), and are increasingly affected by anthropogenic impacts (e.g. habitat modification, fragmentation, climate change; Vörösmarty et al., 2010; Grill et al., 2019). Because of the global impact to freshwater ecosystems, their associated vertebrate populations are declining at alarming rates (83% decline since 1970; WWF, 2018), and their conservation and management are a priority for global biodiversity (IPBES, 2019). Nevertheless, despite broad agreement on the requirements to understand and monitor biodiversity and ecological networks in freshwater habitats (Socolar et al., 2015), our comprehension of biodiversity conservation in this realm lags behind terrestrial and marine environments (Jucker et al., 2018).

The Neotropical region harbours one of the greatest freshwater fish diversities in the world (approximately 30% of all described freshwater fish species), and is currently facing unprecedented levels of anthropogenic pressure. In this region, conservation and management actions in freshwater habitats are challenging due to a lack of infrastructure leading to sampling constraints, as well as a shortage of taxonomic expertise to fully characterise this megadiverse ichthyofauna (Reis et al., 2016). In Neotropical countries, such as Brazil, fish biodiversity assessment relies on sampling using traditional survey methods (e.g. gill nets and traps) followed by morphological identification, which might be selective, harmful, and have low detection rates for rare and elusive species) and small life-stages (Becker et al., 2015; Sales et al., 2018).

Use of specific fishing practices coupled with the remoteness and large geographic extension of most catchments, has meant that Neotropical rivers have not been sufficiently surveyed for baseline estimates of fish diversity. Underestimation of fish diversity resulting from low sampling efficiency may provide biased metrics and hamper management and conservation plans (Trimble & van Aarde, 2012),), including recovery plans for damaged ecosystems (Sales et al., 2018). In addition, with a significantly reduced investment in scientific research and conservation (Thomé and Haddad, 2019), there is an urge to move towards more cost-effective methods to estimate biodiversity at a broad scale (i.e. detecting and monitoring multiple species simultaneously in vast areas).

Molecular approaches offer a universal key to identify, assess and quantify biodiversity, especially in biodiversity-rich and understudied ecosystems and regions (Schwartz et al., 2006). One of the most effective approaches to circumvent the limitations of traditional surveys in mega-diverse systems is the use of DNA barcoding and metabarcoding (Gomes et al., 2015; Cilleros et al., 2019). Sequencing trace DNA present in the water (environmental DNA or eDNA) can now be reliably used to detect species presence (Deiner et al., 2017) and, to some extent, abundance (Doi et al. 2017; Ushio et al. 2018; Shelton et al., 2019). Recently, Cilleros et al. (2019) demonstrated the efficiency of eDNA metabarcoding in providing spatially extensive data on freshwater fish biodiversity in French Guyana, and a better discrimination of assemblage compositions when compared to traditional sampling. We recently showed the influence of sampling medium, as well as sampling preservation and time, on the reconstruction of ichthyofaunal assemblages in a Brazilian catchment, inferred through eDNA (Sales et al., 2019). Nevertheless, the vast majority of eDNA metabarcoding biomonitoring studies remain concentrated in temperate regions, in established and fairly well-accessible environments (Handley et al., 2019; McDevitt et al., 2019).

In this study, we use eDNA metabarcoding to unravel patterns of fish diversity in a poorly studied Brazilian catchment, the Jequitinhonha River Basin (JRB). This catchment belongs to the east Atlantic basin complex, characterised by a high number of species endemism (Reis et al., 2016). Until 2010, the known ichthyofauna of this catchment included 63 described fish species (including 10 introduced species and a substantial number of endangered species, Rosa & Lima 2008; Andrade-Neto, 2010), making this river a relatively low biodiversity ecosystem when compared to its neighbouring basins. This reduced species richness had been linked to historical geological and geographical features (Andrade-Neto, 2010). However, the geological history of the Jequitinhonha is very similar to that of adjacent basins (e.g. Doce and Mucuri river), which led to the consideration that more contemporary factors may explain the low biodiversity in the catchment, including the lack of adequate surveys and impact from anthropogenic activities. The Jequitinhonha is known to be affected by severe droughts, the impact of dams in the main river course and tributaries, and the occurrence of introduced species (Sales et al., 2018;). Thus, an inadequate baseline survey of the basin might still account for a great number of native and cryptic species yet to be described for this catchment (Jerep et al., 2016; Dutra et al., 2016; Nielsen, Pessali & Dutra, 2017).

Furthermore, as other semi-arid and arid regions, the Jequitinhonha faces great variation in water availability (i.e. long dry periods and sudden heavy rain periods; Leite et al., 2010). However, the influence of precipitation in fish assemblages dynamics have not been evaluated in this context. Here we assessed fish diversity, spatially (along the river stem and in two tributaries) and temporally, (before and after heavy precipitation) using eDNA metabarcoding to test whether this DNA-based method can estimate community structure along the course of this anthropogenically-impacted river and thus, be used for biomonitoring purposes.

## 2 MATERIALS AND METHODS

### 2.1 Study Area

The Jequitinhonha River basin (Figure 1), Southeast Brazil (17° S, 43° W), flows between two biodiversity hotspots (‘Cerrado’ and the Atlantic Forest) and is characterised by a tropical climate and environmental heterogeneity. The main river flows over 1,082 km, from its source in Serro, at an elevation of 1200 m, to its outlet in the Atlantic Ocean at the locality of Belmonte. The main river stem is interrupted by two large dams built for hydroelectric power generation: the Irapé, the tallest dam in Brazil, built in 2006, and the Itapebi, established in 2002.

**FIGURE 1.**
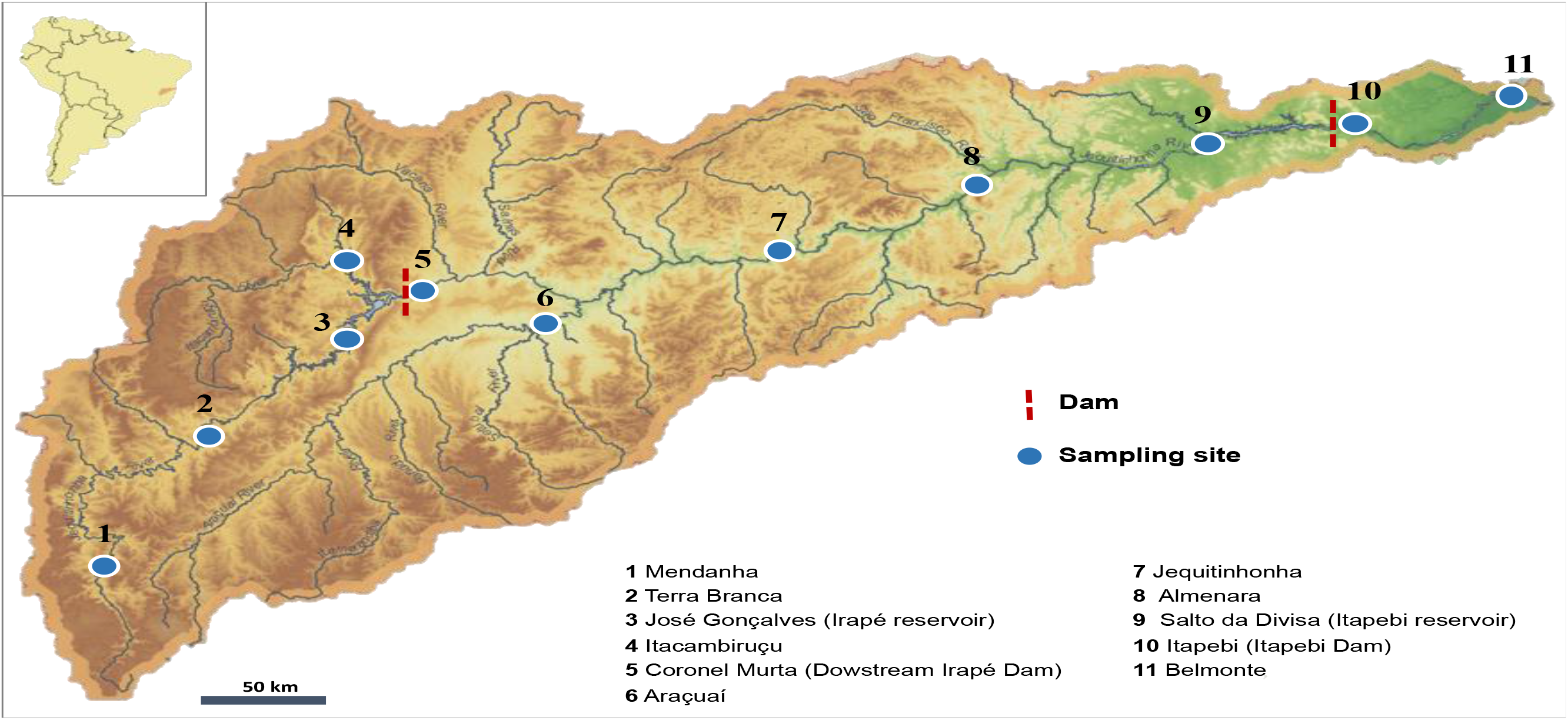
The Jequitinhonha river basin, including sampling sites used in the study, dams and respective hydrological regions.

### 2.2 Historical data and local reference database construction

A compiled species list was built by retrieving all papers available using a Google Scholar search with the terms “fish” and “Jequitinhonha”, combined with a search in Portuguese language journals (applying the terms “peixe”, “Jequitinhonha”, “ictiofauna”), we included data from research papers as well as compiling information on species occurrence from unpublished environmental reports (Table S1, Supplemental information).

To enhance the available reference sequence database in order to obtain a better taxonomic assignment, we retrieved all 12S rRNA mitochondrial gene fish sequences available from GenBank and sequenced 55 additional missing species (Table S2). Information regarding sample preparation and sequencing is provided in the Supplemental information.

### 2.3 eDNA sampling and processing

Two sampling campaigns were conducted at 11 sites during a five-week interval (first sampling period: 22/01 to 01/02/2017; second sampling: 19/02 to 01/03/2017). In between the two sampling campaigns a major precipitation event (from 2.1-50mm in the first sampling event to 100-250 mm in the second sampling event - CPTEC/INPE, 2018) occurred. Sites included locations on the main river (nine) and one on each of two of the major tributaries (the Itacambiruçu river and the Araçuaí river; Fig. 1). At each site, six water samples of one liter each and two sediment samples (^~^25 mL) each were collected. Sediments samples were preserved in ethanol and kept cold during the sampling At the time of sampling proper storage conditions of samples in tropical field conditions had been untested, therefore, we split half of the water samples (N=3) and stored them on ice in a cooling box while for the other samples (N=3) the cationic surfactant benzalkonium chloride (BAC) was added at a final concentration of 0.01% as a preservation buffer to suppress the degradation of DNA by microorganisms (Yamanaka et al. 2017). The effect of storage treatment (ice *vs* BAC) on MOTU diversity recovery was significant only for samples obtained during the first campaign. Still, despite significant (p = 0.016) only 2% of the variance was explained, whereas no significant difference was recovered for samples obtained during the second campaign (Sales et al. 2019), all replicates were used for downstream analyses in this study. In total, 132 water samples and 44 sediment samples were analysed.

Environmental DNA sample filtration, DNA extraction from filtered water and sediment samples, amplification of the 12S rRNA fragment using the MiFish primer set (Miya et al., 2015), multiplexed library preparation, and sequencing of two separate libraries (Library 1/LIB1 – first sampling event; Library 2/LIB2 – second sampling event) in one Illumina MiSeq platform run were conducted as described in Sales et al. (2019), and detailed in the Appendix included in the Supplemental information. Detailed procedures to control for contamination are also described in Supplemental information.

### 2.4 Bioinformatic analyses and taxonomic assignment

The metabarcoding bioinformatics pipeline used for data analysis was based on the OBITools software suite (Boyer et al., 2016), following the protocol described in Sales et al. (2019). Clustering was conducted using a step-by-step aggregation method (SWARM, Mahé et al., 2014) applying a clustering value of d=1 (detailed information on evaluation of different clustering values can be found on Supplemental information). Molecular operational taxonomic units (MOTUs) and the inferred species (based on at at least 97% of similarity with reference sequences; Sales et al., 2020) richness were compared among the three obtained datasets.

For the diversity analyses (species richness and β-diversity), we applied a conservative approach and treated our results as presence/absence-based as suggested by Li et al (2018). Often MOTUs are used as a proxy for species, however, the correlation between these two classifications of diversity are not straightforward. Richness in MOTUs is highly influenced by the occurrence of cryptic species and by the thresholds applied during the bioinformatic analyses (Pawlowski et al., 2018), which may cause an overestimation of true richness (e.g. inflation of different MOTUs belonging to the same species due to natural intraspecific variability, PCR amplification and/or sequencing errors). On the other hand, richness based on MOTUs being assigned to a species may be an underestimate due to the lack of a complete reference database or due to a low taxonomic resolution of the target gene fragment analysed.

To verify whether the inferred community diversity patterns significantly varied because of the species assignment process, two datasets were used for estimating community metrics of alpha and beta. Specifically, the filtered dataset included only MOTUs that could be identified to the rank of species, whereas the non-filtered dataset included all MOTUs retrieved after quality filtering steps. The filtered dataset is a subset of the total MOTU diversity recovered, and thus it provides a more conservative overview for known fish diversity (Li et al., 2018).

A species name assigned to each MOTU might not correspond exactly to the species occurring in the Jequitinhonha River Basin (based on the compiled species list; Table S1) because when the correct species is not present in the reference database, the taxonomic assignment is based on the closest congeneric species. In this case, species not previously reported for this basin are marked with an asterisk in order to highlight that the species herein included might be an indicative of occurrence of the genus and not the exact species present in this river basin.

Statistical analyses were performed in R v3.5.1 (R Core Team 2019). Replicates were pooled (water=6 samples per site, sediment=2 samples) before the following statistical analyses. Alpha-diversity (species richness) was estimated as the total number of MOTUs (unfiltered dataset), or number of MOTUs assigned to species level (filtered dataset), at each sample site. β-diversity was obtained by generating a distance matrix based on the Jaccard coefficient, using the *vegdist* function implemented in vegan 2.5-2 (Oksanen et al. 2013). The Jaccard distance is based on presence or absence of species (value of 0 means both samples share the same species whereas 1 means samples have no species in common). Principal Coordinates Analysis (PCoA) was used to determine the relationship between distance and sites in the β-diversity matrix (cmdscale function) and the correlation between β-diversity and longitudinal distance and the β-diversity and presence of physical barriers (dams) was tested using a Mantel test (Li et al., 2018). The geographic distance matrix between sites was estimated using the road route because the road follows the river course and thus, this distance would provide a better estimate when compared to linear distance between two sample locations. The matrix used for testing the influence of physical barriers was constructed by weighting distance values between sites according to the existence of barriers (e.g. 0 – no physical barrier between sites, 1- one barrier between sites and 2 – two barriers).

Even after our extensive effort to supplement the reference database for taxonomic assignment improvement, most of the MOTUs recovered were not identified to species level (see above) and, thus, a great portion of biodiversity information that could be used for diversity assessments is not included in the reduced filtered dataset. To verify the total diversity recovered and to visualize the community data, we used a hierarchical structure of taxonomic classifications, in the R package Metacoder (Foster et al., 2017). This package, designed for metabarcoding data, provides “heat tree” plots using statistics associated with taxa (e.g. read abundances) and allows for a visual comparison between samples that takes into account their taxonomic/phylogenetic diversity. Venn diagrams were obtained by comparing the orders and families included in the compiled species list, and orders and families detected in each of the eDNA datasets (filtered and non-filtered) using BioVenn (Hulsen, Vlieg, & Alkema, 2008).

## 3 RESULTS

Our extensive review of both published and non-published literature sources resulted in 111 species records for the Jequitinhonha River Basin (Table S1).

We obtained 16.1 million raw reads (LIB1 - 6,399,823; LIB2 - 9,704,699) in one Illumina MiSeq run (See Supplemental information for details). After quality control, clustering and all initial filtering steps, 2056 (LIB1) and 967 (LIB2) MOTUs were kept, with 154 and 59 MOTUs being assigned to species with >0.97 -min-identity, respectively. The number of retained MOTUs varied considerably between filtered and unfiltered datasets and for several species, more than one MOTU was also recovered (Figure 2, Table S3 and Table S4).

**FIGURE 2.**
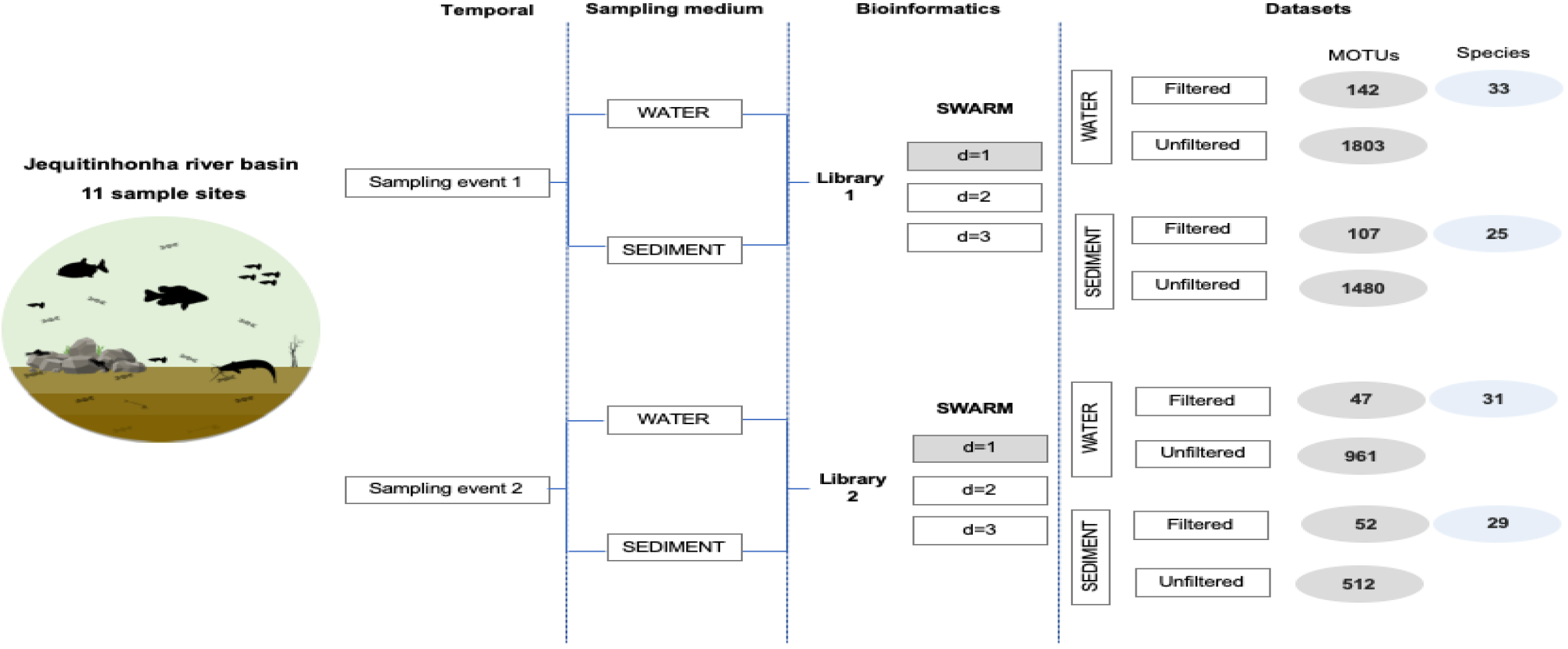
Workflow illustrating the methods used in this paper and respective number of MOTUs retrieved in each dataset analysed, and the final number of species assigned with >0.97 identity.

### 3.1 Taxonomic assignment

Based on the combined data (including all filtered datasets - species >0.97 identity) detected fish diversity included six orders, 20 families, 28 genera and at least 34 fish species (Figure 2, Table S4). Characiformes (n=12) and Siluriformes (n=12) were the two orders represented by the largest number of species identified and all the remaining orders were comprised by less than five species.

A comparison between species identified by eDNA and closely related species reported for the JRB suggests that several congeneric species (e.g. *Leporinus, Prochilodus, Trichomycterus*) are not discernible using our generally applied bioinformatic threshold of 3.0% due to a lack of taxonomically informative variation in the ^~^170 bp fragment of the 12 rRNA gene, for these groups (Table S5).

Comparing the data obtained for both sampling times (Figure 3, Table S6), four species were detected only during the first sampling (*Australoheros facetus, Cyprinus carpio*\*, *Hypostomus* sp., *Trichomycterus* sp.), whilst *Coptodon zilli* and Hoplias intermedius* were detected only in the second sampling.

**FIGURE 3.**
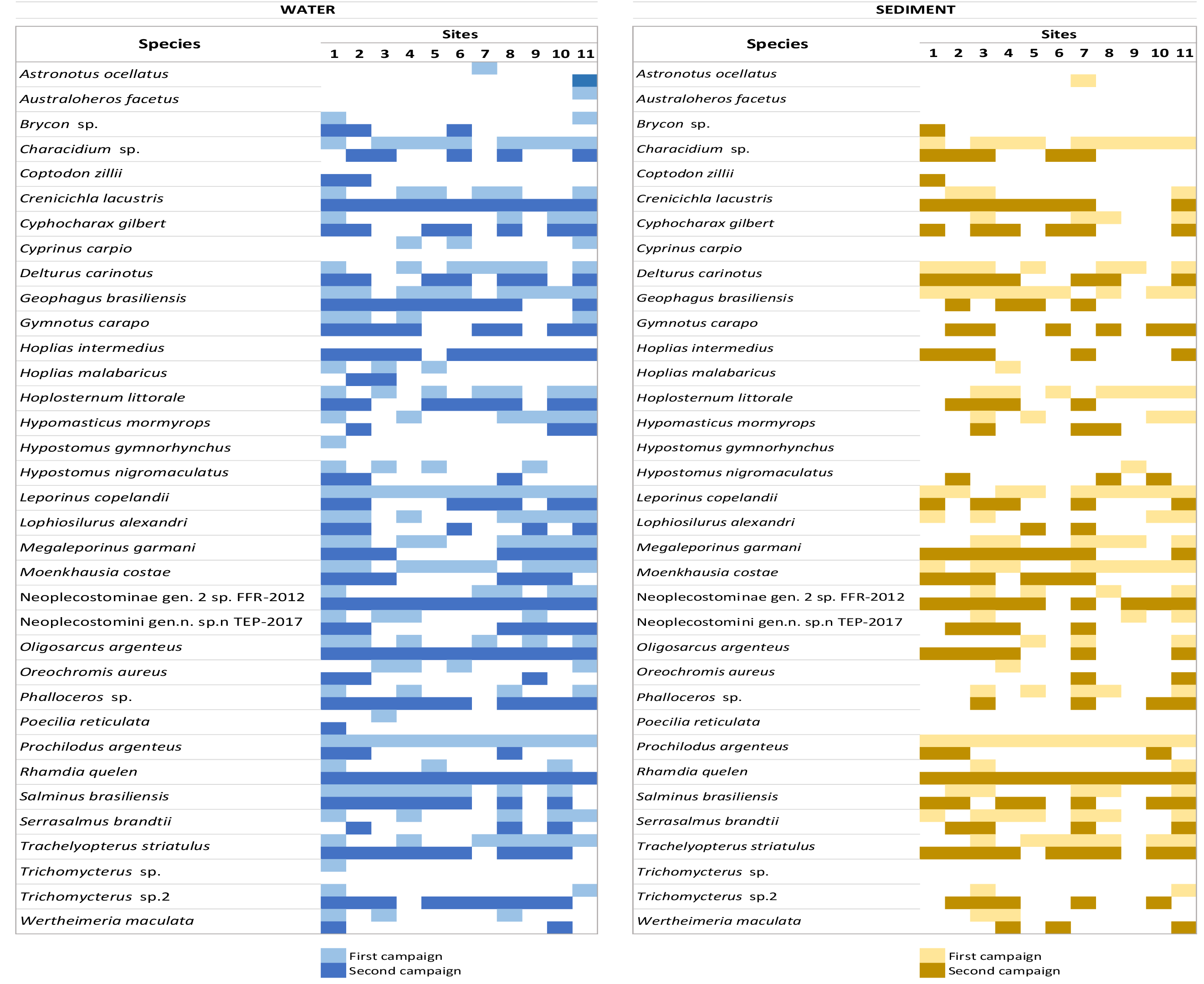
Species distribution in the Jequitinhonha River Basin, according to sampling media and campaign.

Sediment samples failed to detect five species (*Australoheros facetus, Cyprinus carpio*, Hypostomus gymnorhyncus*, Poecilia reticulata, Trichomycterus* sp.), whilst water samples detected all species present in the sediments. Analyses of water and sediment samples demonstrated the occurrence of both widely distributed as well as less abundant species. Several taxa (e.g. *Leporinus* sp., *Prochilodus* sp., *Rhamdia quelen*) were detected in both water and sediment samples in most of sampling sites, in at least one sampling campaign, and therefore seem to have a broad geographic distribution in the Jequitinhonha river basin.

A remarkable result obtained by eDNA included the detection in all analysed sites of species rarely reported in traditional sampling studies (e.g. *Crenicichla* sp., Figure 3). Also we may highlight, the occurrence of putative new records for this basin including invasive species such as the dourado - *Salminus brasiliensis\** and pacamã - *Lophiosilurus alexandri*\*. Furthermore, some species, including native and non-indigenous species, were restricted to a few locations (e.g. native: roncador *Wertheimeria maculata* (sample sites 1, 3, 8 and 10); non-indigenous: oscar *Astronotus ocellatus* (sample site 7); chameleon cichlid *Australoheros facetus* (sample site 11); tilapias *Coptodon* sp.* (sample sites 1 and 2); or were detected in only one campaign (e.g. *Australoheros facetus, Coptodon* sp.*, carp *Cyprinus carpio\**, wolf fish *Hoplias intermedius*, pleco *Hypostomus gymnorhyncus\**, pencil catfish *Trichomycterus* sp.).

The filtered dataset provides a potentially more conservative estimate of fish diversity at the rank of species because many MOTUs could not be assigned a name using the 97% similarity threshold. Fish diversity depicted by the heat trees based on all detected MOTUs (i.e. the unfiltered dataset) shows that diversity remains especially high for the Order Characiformes, as many families appear to be comprised of several MOTUs (e.g. Anostomidae, Prochilodontidae; Figure 4). Comparisons between the filtered and unfiltered datasets demonstrated that a conservative approach (i.e. using filtered data) might lead to a biodiversity information loss since it greatly reduces the diversity in MOTUs recovered and fails in detecting orders and families known to occur in this catchment but that were not identified up to the species level (Figure 5).

**FIGURE 4.**
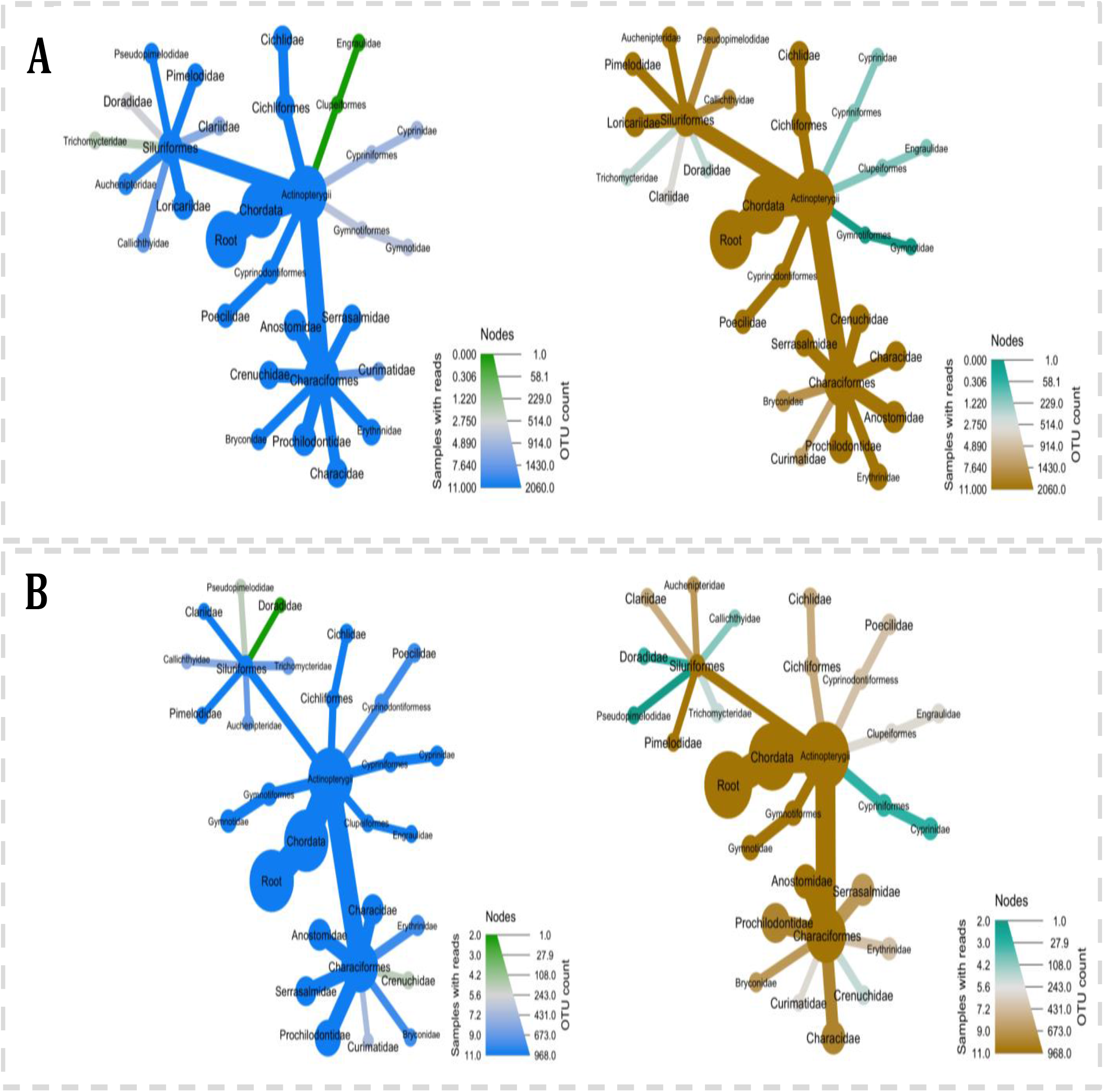
Heat trees displaying the fish diversity recovered for Jequitinhonha River Basin using eDNA metabarcoding unfiltered datasets, during the first (A) and second (B) campaigns. Blue = Water samples; Brown = Sediment samples.

**FIGURE 5.**
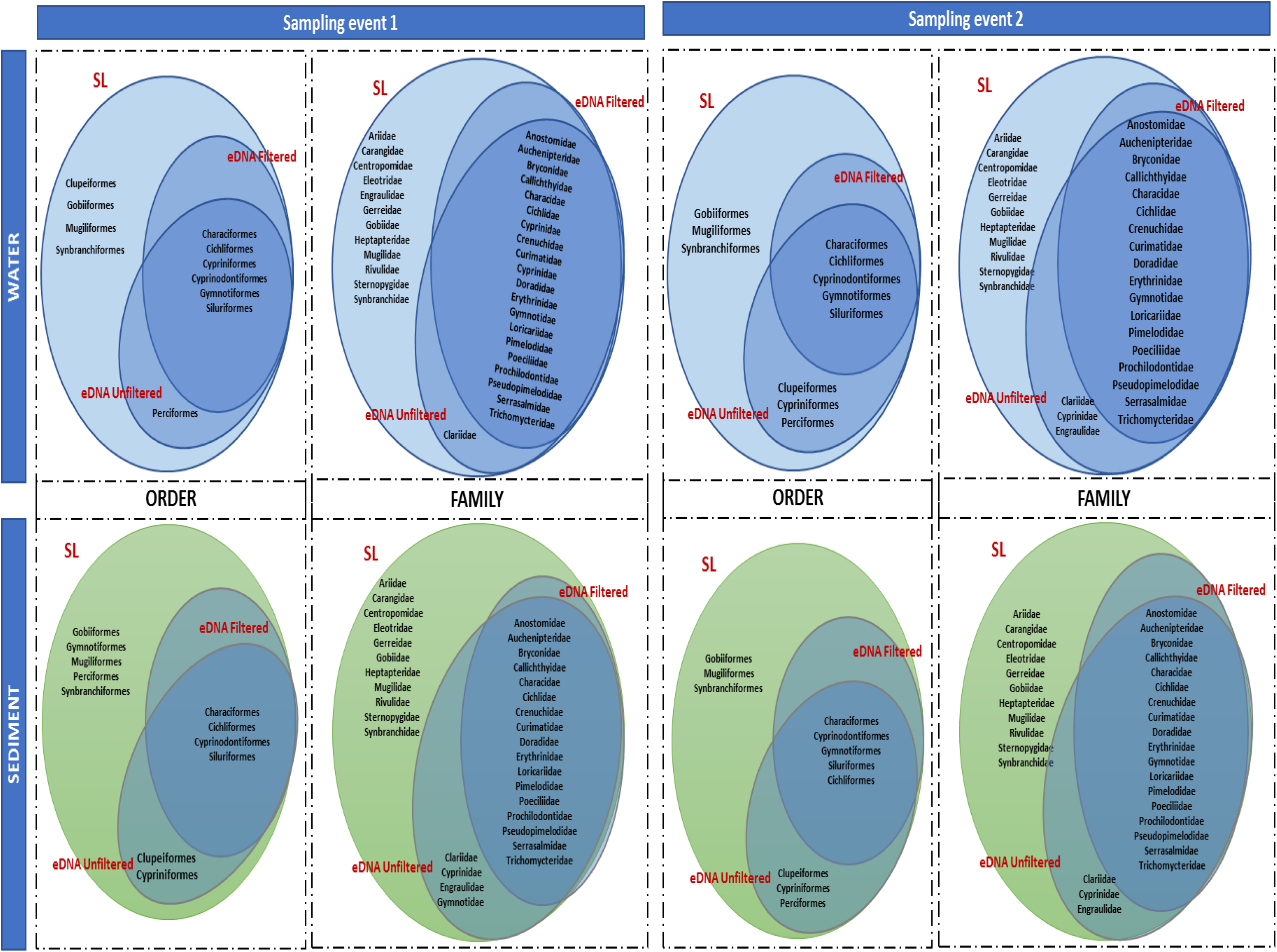
Venn diagram of fish orders and families comparing the data included in the species list based on traditional sampling (SL) to eDNA detected in distinct sampling media (water vs sediment); sampling campaign; and datasets analysed (unfiltered vs filtered).

### 3.2 Species richness and Beta diversity

During the first campaign, highest MOTU richness was found in water samples from the most upstream (site 1) and downstream (site 11) sampling sites, followed by sampling sites 4 and 8 (Figure 6A). The lowest number of MOTUs was recovered for sample site 7. Beta diversity patterns showed similarities between sample sites 4 and 11, and sample sites 1 and 8, whereas sample site 7 showed the most distinct fish assemblage when compared to all locations. Environmental DNA recovered from water samples collected three weeks later, demonstrated that species richness among sites fluctuate in time in this catchment (Figure 6B), with generally greater homogeneity in the species richness amongst all sample sites in the late sampling event. Still, the most upstream and downstream locations (1, 2, 10, 11), alongside sample site 8, still harboured the highest number of species.

**FIGURE 6.**
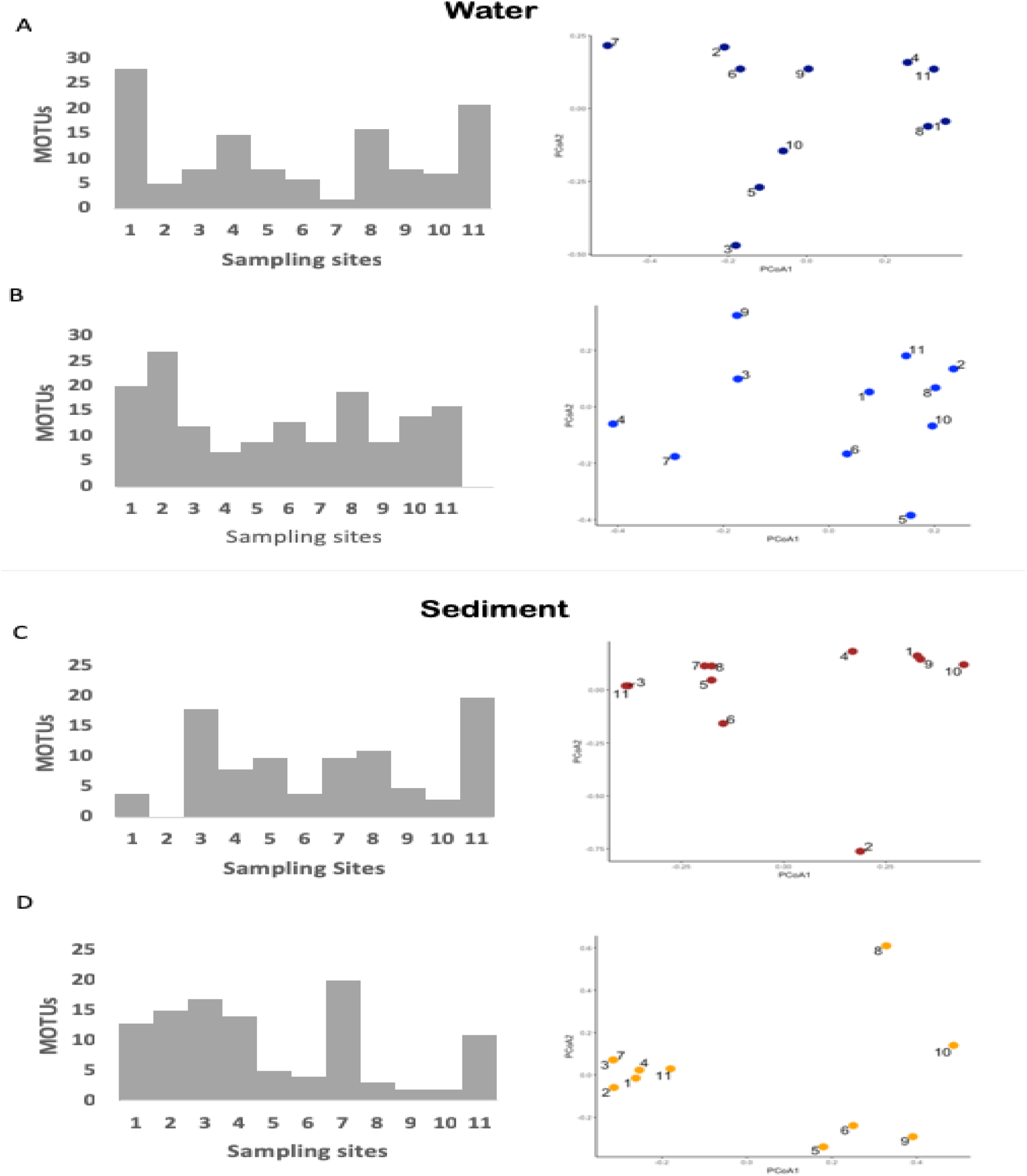
Filtered dataset, showing the species richness distribution along the Jequitinhonha River Basin and Principal Coordinates Analysis (PCoA) of β-diversity of sampling locations (Jaccard distance). A) Water samples obtained in the first campaign; B) Water samples obtained in the second campaign; C) Sediment samples obtained in the first campaign; D) Sediment samples obtained in the second campaign.

Data recovered from sediment samples provided a different overview of species richness and beta diversity. Overall, the number of species recorded for sediment samples was lower compared to water samples in the first campaign (Figure 6C). Sample site 1 had a much lower species richness compared to water samples along with sampling sites 2, 4, 8, 9, 10. An increase in the species richness was detected for sampling sites 3, 5 and 7, while sample sites 11 and 8 were confirmed as highly species-rich locations. In the second campaign (Figure 6D), when compared to data recovered from water samples, six sample sites (1, 2, 6, 8, 9, 10) had a lower species richness, while higher values were obtained for sample sites 3, 4, 7.

Over time, the pattern of harbouring the highest species richness appeared relatively constant in sites 1 and 11 for both sampling media, except in the first campaign where fewer species were detected in location 1 for sediment. Yet, the most downstream location kept an almost stable species richness in both sampling media for both sampling campaigns.

Longitudinal distance had a negligible effect on beta diversity amongst sample sites (*p-* value > 0.05, Table 1) and the presence of physical barriers (e.g. dams) also did not show a significant influence on beta diversity of different sample types (water and sediment, Table 1). A positive significant correlation was found between filtered and unfiltered datasets, for both water and sediment (Table 1.)

**TABLE 1.**
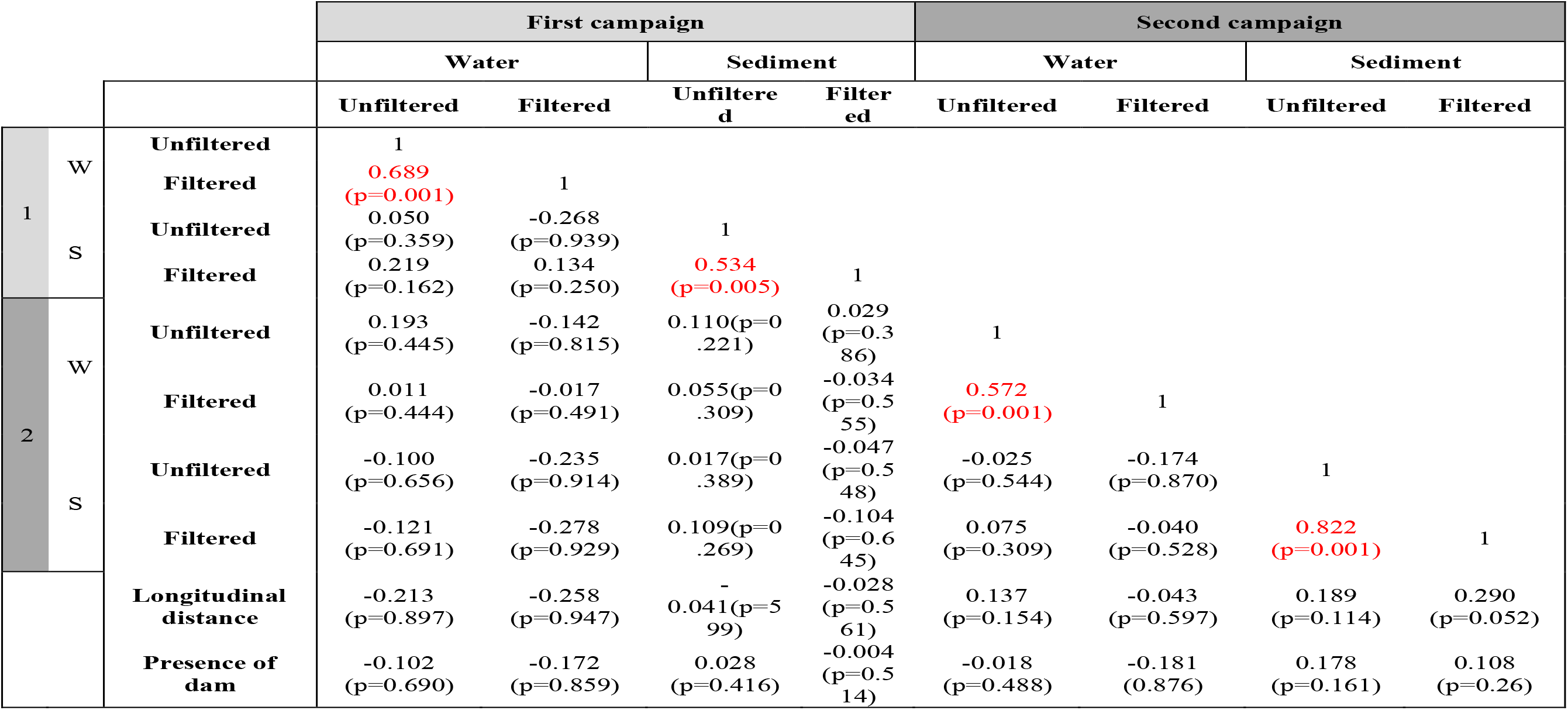
Mantel *r* and p-values (in parentheses) for all the pairwise comparisons between datasets, sampling media, geographic distance and presence of barriers (dams).

For both sampling media, despite the variation in taxa richness showed by both datasets, the pattern of alpha diversity variation among sample sites obtained for filtered (species) and unfiltered (MOTUs) datasets were still quite congruent (Figure 7). However, for sediment samples collected in the first campaign, sites 3 and 11 had a greater MOTU diversity when compared to all nine remaining locations (Figure 7C). Despite also being the most species rich sites, the great amount of MOTUs obtained and not assigned indicates that a great diversity remains hidden in this sampling medium. Also, as demonstrated by the PCoA (Figure 7C), in the first campaign these sites had a more distinct fish assemblage when compared to the others. Furthermore, a higher resolution was obtained for the unfiltered dataset as a more segregated sample clustering is evident in the PCoA ordination. Sediment samples from the first campaign exhibited a peculiar clustering, with highly diverse samples in 3 and 11 strongly separated from all other sites.

**FIGURE 7.**
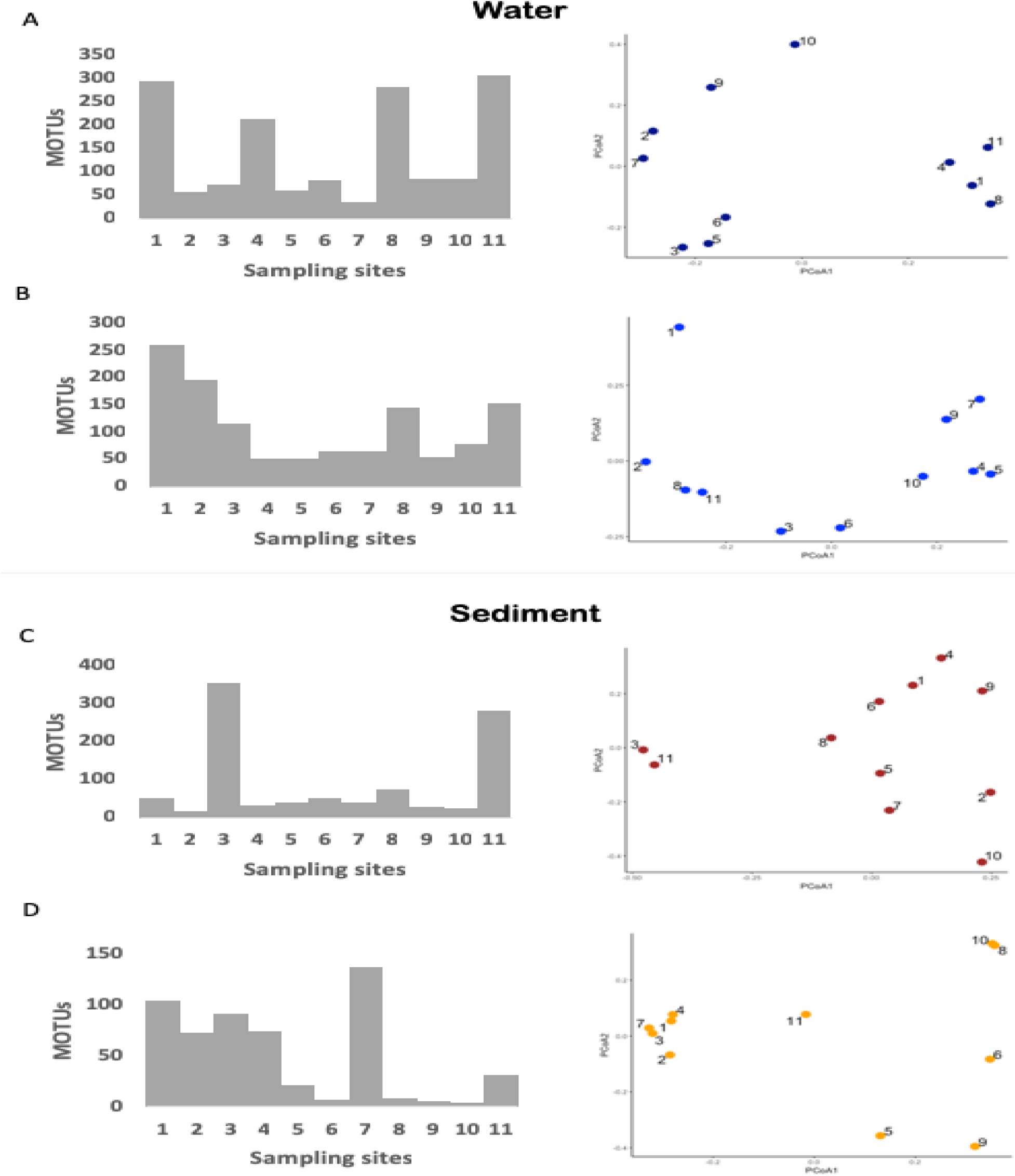
Unfiltered dataset, showing the species richness distribution along the Jequitinhonha River Basin and Principal Coordinates Analysis (PCoA) of β-diversity of sampling locations (Jaccard distance). A) Water samples obtained in the first campaign; B) Water samples obtained in the second campaign; C) Sediment samples obtained in the first campaign; D) Sediment samples obtained in the second campaign.

### 4 DISCUSSION

The understanding of species distribution and the processes shaping spatial variation and community composition are crucial for applying sustainable management schemes and ensure timely conservation of biodiversity, especially for endemic and threatened species. Such actions also require methods that allow for rapid and robust detection of biodiversity at different spatial scales (Kelly et al., 2014). Here, we used eDNA metabarcoding of water and sediment samples to investigate fish community variation over time along the course of a Neotropical river.

We found that eDNA metabarcoding applied to understanding fish distributions in a neotropical setting greatly enhanced our ability to not only measure richness along the course of a large river, but also to reveal hidden diversity and putative unrecorded species invasions. The compiled list of species (N=111) reported for the Jequitinhonha river basin herein was higher than previously recorded (N=63) in 2010 (Andrade-Neto, 2010). Hence, our thorough evaluation of all possible taxonomic information available at the time of our study estimates the occurrence of more than 80 species in this catchment (Andrade-Neto, 2010; Godinho et al., 1999). Our molecular assessment based on eDNA metabarcoding demonstrates that, as of yet, there may be even more species yet to be recorded and putting the richness of this basin on par with other closely adjacent basing thought to harbour higher diversity. These results demonstrate our current lack of understanding of tropic diversity in many systems and corroborates that new DNA based methods are ideal in generating new baselines for biodiversity monitoring.

### 4.1 Introduced and native species

Environmental DNA metabarcoding allows the detection of multiple species simultaneously, including species not expected to occur in an area (Deiner et al., 2017), helping to track biological invasions and providing an early warning of species introduction. Here, almost 30% of the taxa detected by eDNA were non-indigenous species, including species not reported yet for this catchment. To our knowledge, previous records of *Salminus brasiliensis* and *Lophiosilurus alexandri* occurrence in the JRB are absent from the literature. These are commercially important species, already introduced for fishery purposes in several Brazilian basins (Vitule et al., 2014). Hence, their occurrence in the Jequitinhonha is not necessarily a surprise. However, it raises concerns about the ecological consequences of such unmanaged introductions. Biodiversity loss is not only restricted by species disappearance, but also by a reduction in ecosystem services due to an increase of biological similarity between areas (i.e. species loss or increase through biological introductions leading to biotic homogenization; Rahel, 2000).

It has been widely documented that analysis of eDNA surpasses traditional methods for assessment of biodiversity and detection of invasive species (Schmelzle & Kinziger, 2016; McDevitt et al., 2019). The only cyprinid previously documented in this basin was *Hypophthalmichthys molitrix*. Herein, we registered the presence of *Cyprinus carpio*, another species that has been widely introduced to Brazilian waters (Alves et al., 2007). Environmental DNA metabarcoding also detected various species of tilapia (*Oreochromis* sp. and *Coptodon zilli*). The impacts of tilapia invasion are well known worldwide, and all species show high invasive potential, including in Neotropical countries (Cassemiro et al., 2017).

Our study also detected remarkable cases, such as the native species *Crenicichla* sp. The genus *Crenicichla* is one of the most species rich among the South American Cichlids, where it is known to widely occur. However, the genus is still lacking an improved taxonomic resolution and conservation status evaluation (Kullander & de Lucena, 2006). In 2006, an expedition applied extensive sampling efforts to collect *Crenicichla* sp. in the Jequitinhonha, without any success, and this species was only documented in 2009 by an environmental report based on traditional sampling and morphological identification (Kullander & Lucena, 2006; Intertechne, 2009). An issue reported worldwide, is that even when monitoring programmes are conducted, most of the data obtained are often not published or made available and thus remain inaccessible to further scientific studies (Lindenmayer & Likens, 2009; Revenga et al., 2005). Here, eDNA metabarcoding data revealed that this species might be present at several locations in the Jequitinhonha, indicating a possible large geographical distribution.

Taxonomic issues are often present in monitoring programs and the risk of misidentification exists, regardless of the method applied (i.e. traditional sampling, morphological identification, eDNA; Radinger et al., 2018; Jerde, 2019). Erroneous identifications might also be present in the reference databases, especially in highly biodiverse regions such as the Neotropics, where the amount of unknown and undescribed taxa and the occurrence of cryptic species represent substantial issues. As demonstrated in previous studies, identification of some species might be problematic when using eDNA metabarcoding based on the 12S fragment employed here, due to its lack of taxonomic resolution and the incompleteness of the reference databases (Yu et al., 2012; Eiler et al., 2013). Because a gene tree is not necessarily related to a species tree, the phylogenetic resolution it provides can by obscured for groups of taxa. The imperfect taxonomic resolution might allow the multiple assignment of congeneric species (i.e. one species being concomitantly assigned to its multiple congeners) when several reference sequences are available (please see example of *Prochilodus* sp. below). In contrast, when the reference database is not complete for all species occurring in the area, several MOTUs belonging to distinct species might be assigned to and errouneously identified as the single closely related species available in the database (Sales et al., 2020) For instance, most MOTUs belonging to *Prochilodus* sp. could not be assigned to species level due to a high similarity among orthologous sequences from congeneric species. This poses a conservation issue, since *Prochilodus argenteus* is an invasive species in the Jequitinhonha, and is believed to have recently diverged from the endemic species *P. hartii* (Melo et al., 2018). Henceforth, due to the conservative criteria applied to analyse the data, the number of species detected is surely underestimated.

Six anostomids are described for the Jequitinhonha, and here we identified one of these species (*Megaleporinus garmanii*), but also identified two species not previously reported (*Leporinus copelandii* and *Hypomasticus mormyrops*). The only previous record of *Leporinus copelandii* was deemed as an historical error (Andrade-Neto, 2010). Cilleros et al. (2019), despite using a different 12S fragment, also reported the limitations in the taxonomic assignment of species belonging to the genus *Leporinus*, therefore our data set is unable to clarify the nuances within this group.

### 4.2 Anthropogenic impacts and species richness

Ecological communities vary in time and space, and the monitoring of these dynamics is essential for conservation purposes (Bálint et al., 2018). In the Jequitinhonha River basin, significant spatial and temporal fluctuations in fish assemblages inferred from eDNA were detected. The longitudinal distance and presence of barriers did not explain community variation (p>0.05); however, anthropogenic impacts might still have an influence on fish diversity distribution in this river basin. Regarding data recovered from water samples, low species richness were recovered from the reservoirs (3 – José Gonçalves, 9 – Salto da Divisa) and the first sites located downstream the dams (5 – Coronel Murta and 10 – Itapebi). The presence of dams is a well known fish diversity reduction factor since these barriers greatly impact the environment (i.e. modification of physical and ecological characteristics of the habitats, such as modifications in water flow, nutrient dynamics, water quality and temperature; Pelicice & Agostinho, 2007; Pompeu et al., 2012). However, changes in fish distribution and communities composition may also arise from plenty of distinct alterations and complex interactions in the impounded environment (Agostinho, Pelicice & Gomes, 2008). Therefore, despite no significant correlation between dams and fish diversity was herein found, the use of eDNA metabarcoding offers a promising tool for evaluating the impoudment’s impact on fish distribution and thus, should be further investigated.

The sites comprising the highest fish diversity in this basin were represented by locations characterized by different anthropogenic influences. The most upstream site (Mendanha) is located in a less populated and more pristine region (Table S7, Supplementary Material), near two areas of natural preservation (State Parks Biribiri and Rio Preto). The other two sampling sites (Almenara, 8, and Belmonte, 11) are located near more densely populated cities and impacted areas (i.e. due to the deforestation and mining activities, siltation increases towards the river mouth and represents one of the greatest impacts in the Jequitinhonha river - IBGE, 1997). Almenara, is a particularly impacted area, and during the sampling had a low water level and accumulation of sediments, which might have contributed to increase the eDNA concentration and accumulation, increasing the species diversity, despite the low environmental quality.

The putative effect of anthropogenic activities on fish eDNA recovery herein described corroborates the well-known impacts of human actions (e.g. construction of dams, species introduction, pollution) leading to biotic homogenization (Agostinho et al., 2008, Ribeiro et al., 2017).

### 4.3 Seasonal changes in fish assemblages

Seasonal changes driven by natural factors (e.g. water flow, rainfall) could also contribute to explain assemblage variation even over a short time frame (i.e. weeks) as mobile species, such as fish, can rapidly disperse and vary their distribution in response to changing abiotic conditions (Arrington & Winemiller, 2006; Fitzgerald et al., 2017).

Water availability shows a great temporal variability in semi-arid and arid regions, with short, but intense, rainfall episodes followed by long dry periods (Leite et al., 2010). The Jequitinhonha river basin is inserted in a semi-arid region and in the first sampling campaign it was facing a severe drought. Before the second sampling campaign, an increase in the average accumulated rainfall (from 2.1-50mm in the first sampling event to 100-250 mm in the second sampling event; CPTEC/INPE, 2018) might have contributed to a higher evenness in MOTU richness/fish diversity amongst sample sites (regarding the contemporary species richness inferred through water samples). The climatic and hydrological changes followed by the onset of the rainy season usually triggers the start of fish migration in the semi-arid regions (Chellappa et al., 2003; Chellappa et al., 2009). An increased water volume and subsequently higher connectivity of aquatic habitats might stimulate the dispersal and result in reduced densities of organisms (Fitzgerald et al., 2017). Therefore, the result here presented might suggest that freshwater fish assemblages in tropical habitats may vary significantly between dry and wet seasons. Besides the apparent homogenization found after the rainfall event, an important factor to take into consideration is the reduction of diversity recovered in the second campaign when compared to the first. The ecology of DNA might play an important role regarding this matter, as eDNA molecules could be more diluted in the water column decreasing the detectability of some species (e.g. rare or less abundant species).

### 4.4 eDNA transport and species richness

Another factor we need to take into account is eDNA transport from locations upstream from our sample sites. This transport could lead to an overestimation of species richness recovered for each sample site, and, the species identification per site therefore does not mean that the species themselves are present there at the time of collection (Barnes & Turner, 2016; Deiner et al., 2014). Still, eDNA transport distances may vary between river systems due to abiotic and biotic factors (e.g. temperature, pH, bacterial load, or seasonal changes such as drought or intense rainfall periods; Deiner et al., 2016). Most of the studies evaluating the effect of eDNA upstream transportation reported travel distances of few kilometers, whereas, a travel distance higher than 100km was demonstrated by Pont et al. (2018) for a high discharge (m3/s) river system. Still, despite the eDNA downstream transportation, the latter study demonstrated the capability of eDNA in providing an accurate snapshot of fish assemblage composition in a large river and finally, suggested that a distance of around 70 km would be enough to limit the potential noise of eDNA transport. Therefore, despite having a high discharge rate (average of 409 m^3^/s), the approximate distance between sites was 100 km and thus, the influence of eDNA transport on species detected at each site might not be considered as a great concern here. However, as no study has been conducted in Brazilian lotic environments focusing on understanding eDNA transport and diffusion, it is difficult to draw sound conclusions regarding this matter and so, additional studies focusing on the information recovered from eDNA in large neotropical rivers might contribute to expand the knowledge of its complex spatiotemporal dynamics.

The high alpha diversity values found for the site located at the river mouth (site 11, Belmonte) deserves some consideration since this region has marine influence (including the detection of one marine family, Engraulidae, by sediment samples in this sample site, Figure 4) and its abiotic characteristics (e.g. increased salinity) would be expected to restrict the occurrence of some freshwater species. A hypothesis that could explain the detection of species not expected to occur in this area includes eDNA transport and accumulation. Species shed DNA constantly, which can be available in the water column or bound to superficial sediment. A higher concentration and longer persistence of fish eDNA in the sediments might contribute to eDNA molecule resuspension which might affect inferences from aqueous DNA in both spatial and temporal scales (Turner et al., 2015; Graf & Rosenberg, 1997; Bloesch, 1995;).

Due to the fragmentation of the Jequitinhonha River, this site (site 11, Belmonte) is located in a region characterized by a high level of sediment trapping (freeflowingriver.org/maptool/) and possibly, this segment can act as an “eDNA reservoir” due to the accumulation of molecules transported throughout the river. In addition to that, an increase in water flow and tidal movements can also cause eDNA particle resuspension (increasing the probability of retrieving old eDNA from the sediment beds – Jamieson et al., 2005), which, associated with the resistance applied by the incursion of the marine waters into the river, can contribute to retain and resuspend the eDNA accumulated in this area, making it available in the water column. Considering this, river mouths should then be further investigated as putative eDNA reservoirs since it could contribute in future sampling strategies focusing on obtaining a snapshot of the entire fish community at a large scale.

Bioinformatics and technical aspects also play an important role in diversity recovery from eDNA samples, and the existing trade-off between uncertainty and stringency may be carefully considered when interpreting eDNA results as it might lead to false negative or false positive detections (Evans et al., 2017; Grey et al., 2018). Regarding the analysed datasets, the filtered data is considered as a subset of the total diversity recovered and showed a lower diversity at the order and family levels. However, the significant positive correlation between datasets demonstrated that beta-diversity is not influenced by the filtering criteria applied as much as the effect of sampling medium or sampling time. As suggested by Li et al. (2018), the filtered dataset provided a more conservative overview of fish diversity, compared to the unfiltered dataset and thus did not detect several families and orders known to be present in this catchment.

Fish diversity depicted by the heat trees based on the unfiltered data shows that a hidden diversity might be present, especially for the Order Characiformes, as many families appear to comprise several MOTUs (e.g. Anostomidae, Prochilodontidae). This likely reflects the presence of multiple genera/species such as in the Anostomidae, known to harbour at least seven species in this basin, which are absent from the reference sequence databases. Therefore, to avoid underestimating the biodiversity, and reduce ambiguity in eDNA-based species detection, we stress the importance of coordinating morphological surveys alongside DNA assessments. Most importantly, there is also a need of increasing efforts towards building more complete genetic reference databases, ideally composed of whole mitochondrial genomes, as the lack of reference sequences has been considered as a great hindrance to fullfill the potential of eDNA metabarcoding in assessing biodiversity rich ecosystems (Cilleros et al., 2019; Sales et al., 2020).

Given the unprecedented rates of population and species decline and the increasing anthropogenic impacts on freshwater communities, the importance of a rapid, robust and efficient monitoring program has never been more in need for this ecosystem. Here we illustrated eDNA ecology when analysing an entire river basin from the headwater to the river mouth, and highlighted some of the challenges of applying eDNA metabarcoding in spatio-temporal ecological studies, including recommendations for future work. Understanding eDNA metabarcoding dynamics is an important step to make it a complementary monitoring tool to traditional methods. This enhancement can improve the applicability of eDNA metabarcoding for biomonitoring purposes in Brazilian freshwaters and therefore, allow the detection of elusive, rare or patchily distributed species and provide data for neglected and difficult to access localities.

## ACKNOWLEDGEMENTS

This work was supported by Conselho Nacional de Desenvolvimento Científico e Tecnológico (CNPq) and Science without Borders Program (Grant #204620/2014-7). NGS also thanks FCT/MCTES for the financial support to CESAM (UID/AMB/50017/2019), through national funds. DCC is grateful to CNPq for the productivity fellowship (CNPq 306155/2018-4). We are grateful to Letícia Sales for the invaluable fieldwork assistance and Gilberto Nepomuceno Salvador for assistance with the map.

## DATA ACESSIBILITY

Raw data will be made available on DRYAD upon acceptance.

## AUTHOR CONTRIBUTIONS

NGS, OSW and SM designed the study. NGS carried out the fieldwork. NGS and OSW performed the laboratory work and the bioinformatics. NGS analysed the data primarily, with contributions from ADM, IC, KD and KP. All authors discussed the results and implications. NGS drafted the manuscript, all authors provided manuscript input and contributed in discussion that developed the study.

