## Supporting information for "Space-time dynamics in monitoring neotropical fish communities using eDNA metabarcoding"

**Table of Contents:**


| | |
|---|---|
| **Appendix 1.** Elaborated Material and Methods | Page 2 |
| **Table S1:** Species reported for the Jequitinhonha River Basin | Page 7 |
| **Table S2:** List of samples sequenced for the custom reference database. | Page 10 |
| **Table S3**: Comparison of different SWARM clustering thresholds. | Page 11 |
| **Table S4**: Species identified applying the minimum identity of 0.97, according to each SWARM threshold. | Page 12 |
| **Table S5:** Taxa detected by eDNA metabarcoding and correspondent nearest neighbour species reported for Jequitinhonha river basin | Page 13 |
| **Table S6**: Taxa detected in each sampling event and sampling medium | Page 14 |
| **Table S7**: Sample sites including code, city, human population* and GPS coordinates. | Page 15 |

### APPENDIX

#### 1 Elaborated Materials and Methods

*1.1 Sequencing of tissue DNA*

Tissue samples of 108 specimens belonging to 55 neotropical fish species were obtained from the Laboratório de Genética da Conservação tissue collection (LGC), at Pontifícia Universidade Católica de Minas Gerais (PUC Minas). DNA was extracted from fin clips, using DNeasy Animal tissue DNA extraction kit (Qiagen) following the manufacture's protocol. Fragments of the mitochondrial 12S rRNA gene were amplified using the MiFish primers (Miya et al., 2015) and the polymerase chain reaction (PCR) consisted of: 1.0µl of buffer (10x) including $MgCl_2$(50mM,) 0.3µl of dNTP (total of 10mM), 0.25µl of each MiFish primer (10µM), 0.2µl of BIOTAQ DNA polymerase (5U/µl) (Bioline), 7.0µl of ultrapure water, and 1.0µl of DNA template (10 ng/µl). PCR conditions consisted of an initial step of 10 min at 95 °C followed by 35 cycles of 30s at 95 °C, 45s at 60°C, and 30s at 72°C and one final step of 5 min at 72 °C. PCR products were visualised on 1% agarose gels and successfully amplified samples were sequenced by Macrogen Laboratories (www.macrogen.com) or included in one MiSeq run (V2 nano kit, 2x150bp) following the same procedures for library preparation and analysis as described for the eDNA samples.

*1.2 eDNA sampling and processing*

In total, 132 (1 Litre) water samples and 44 sediment samples were collected for two times point across 11 sites. Water was filtered approximately 8 hours after collection, using mixed cellulose esters (MCE) filters (diameter: 47 mm, pore size: 0.45 mm, Merck Millipore) using an automatic vacuum pump. Filters were removed, folded and stored at −20°C in 15 mL tubes containing silica beads (Majaneva et al., 2018). Sediment samples (˜25 mL) were preserved on site by storing them in 50 mL centrifuge tubes and adding in 100% ethanol (˜25 mL).

DNA extraction from the filters was conducted using the DNeasy PowerWater Kit (Qiagen) and DNA from the sediments was extracted using DNeasy PowerMax Soil Kit (Qiagen), following the manufacturer's protocol in both cases. DNA concentration was determined in a Qubit fluorometer (Qubit dsDNA HS Assay Kit, Invitrogen).

*1.3 Amplification, library preparation and sequencing*

Amplicons of 169-172bp from a variable region of the mitochondrial 12S rRNA gene were obtained with the MiFish primers (MiFish-U-F, 5′- GCCGGTAAAACTCGTGCCAGC-3′; MiFish-U-R, 5′- ACATTATCATAGTGGGGTATCTAATCCCAGTTTG -3′, Miya et al., 2015).

Samples (N= 176) and negative controls (N=7) were sequenced in a single multiplexed Illumina MiSeq run, along with 54 additional samples belonging to a non-related project (not included in this study). For the present study, two libraries were sequenced, containing a total of 183 samples including collection blanks (N=3) and laboratory negative controls (N=4), using two sets of 96 primers with seven-base sample-specific oligo-tags and a variable number (2-4) of leading Ns (fully degenerate positions) to increase variability in amplicon sequences. PCR amplification was conducted using a single-step protocol and to minimize bias in individual reactions, PCRs were replicated three times for each sample. The PCR reaction consisted of a total volume of 20 µL including 10 µl AmpliTaq Gold™ 360 Master Mix (1X; Applied Biosystems); 0.16 µl of BSA (5mg); 1 µl of each of the two primers (5 µM); 5.84 µl of ultra-pure water and 2 µl of eDNA template. The PCR profile included an initial denaturing step of 95°C for 10 min, 40 cycles of 95°C for 30s, 60°C for 45s, and 72°C for 30s and a final extension step of 72°C for 5 min. Amplifications were determined by electrophoresis in a 1.5% agarose gel stained with GelRed (Cambridge Bioscience). PCR products were pooled in two different sets and purified using MinElute columns (Qiagen), and Illumina libraries were built from each set, using a NextFlex PCR-free library preparation kit (Bioo Scientific). Size selection was performed using 1.1x Agencourt AMPure XP (Beckman Coulter), libraries were then quantified by qPCR using a NEBNext qPCR quantification kit (New England Biolabs) following the manufacturer's protocol and pooled in equimolar concentrations along with 1% PhiX (v3, Illumina). The libraries were run at a final molarity of 10pM on an Illumina MiSeq platform in a single MiSeq flow cell using the 2x 150bp v2 chemistry.

*1.4 Bioinformatic analyses*

The bioinformatic analyses were conducted using the OBITools metabarcoding package (Boyer et al. 2016). High values of fish biodiversity expected in Neotropical realms and scarcity of reference sequences pose additional challenges to delimiting taxonomic units and assignment of their representative sequences. Therefore, we used a step-by-step aggregation method (SWARM, Mahé et al., 2014) and evaluated the impact of different values for clustering (i.e. d value) on species recovery. Thus, we generated three datasets to evaluate the influence of using different values for the clustering distance d ($d$=1, $d$=2, and $d$=3) and the value of d=1 was chosen for further analysis since higher values produced a reduced recovery of species (Table S3). The taxonomic assignment was performed using ecotag with a custom reference database built retrieving all 12S rRNA sequences available from GenBank (access date: May 2018) and the local reference database built in this study. MOTUs not matching Actinopterygii were removed, each MOTU was assigned to species based on a 97% sequence similarity to references (identity 0.97). The *cut off* was established based on published eDNA metabarcoding studies which applied this threshold (>97%) for detecting a great variety of fish species (Li et al., 2018; Nakagawa et al., 2018); MOTUs well represented but showing <97% similarity to references were presented and discussed as putatively belonging to species still absent in the database.

### 1.5 Contamination control procedures

Contamination control procedures were implemented both in the field and in the laboratory. For collection of eDNA samples we used disposable sterile collection bottles, disposable gloves, and pre-treated all equipment and surfaces with 50% bleach solution for 10 minutes, followed by rinsing in distilled water before and after each use. Filtration blanks were created between the processing of replicates for each sample site and time to test for potential contamination during the filtration stage and to monitor the risk of false positives due to cross sample contamination during laboratory processing. Field blanks were obtained through filtering clean water, following the same protocols and processed and preserved exactly in the same way as field samples. To account for possible contaminations arising during the laboratory work, non-template amplification controls (NTC, i.e. several reactions lacking DNA template) were included during the amplification step (PCR) alongside with all other samples, and processed using the same exact following protocols.

Tag jumping is a challenge for pooled samples analysed on the Illumina sequencing platform (Schnell et al, 2015; Olds et al. 2015), to account for this issue, we adopted a stringent approach to guarantee the removal of false positives and MOTUs putatively originated by sequencing error or contamination. Thus, for each MOTU the total number of reads detected in the negative controls (corresponding to this MOTU) were subtracted from all samples, then MOTUs containing less than 5 reads in total were removed before analysis.

Table S1: Species reported for the Jequitinhonha River Basin.

| Order | Family | Species | Status | Habitat | Reference | Current status (Catalog of Fishes) |
|---|---|---|---|---|---|---|
| Characiformes | Anostomidae | Hypomaticus garmani | native | freshwater | Andrade-Neto, 2009 | Megaleporinus garmani |
| Characiformes | Anostomidae | Leporinus bahiensis | native | freshwater | Andrade-Neto, 2009 | Leporinus bahiensis |
| Characiformes | Anostomidae | Leporinus crassilabris | native | freshwater | Andrade-Neto, 2009 | Synonym of Megaleporinus elongatus |
| Characiformes | Anostomidae | Leporinus elongatus | | freshwater | Pugedo et al., 2016 | Megaleporinus elongatus |
| Characiformes | Anostomidae | Leporinus garmani | native | freshwater | Report CEMIG, 2005 | Megaleporinus garmani |
| Characiformes | Anostomidae | Leporinus spp. | not described yet | freshwater | Andrade-Neto, 2009 | |
| Characiformes | Anostomidae | Leporinus steindachneri | native | freshwater | Andrade-Neto, 2009 | Leporinus steindachneri |
| Characiformes | Anostomidae | Leporinus taeniatus | | freshwater | Pugedo et al., 2016 | Leporinus taeniatus |
| Characiformes | Bryconidae | Brycon devillei | endangered | freshwater | Andrade-Neto, 2009 | Brycon devillei |
| Characiformes | Bryconidae | Brycon sp. | not described yet | freshwater | Andrade-Neto, 2009 | - |
| Characiformes | Callichthyidae | Callichthys callichthys | native | freshwater | Andrade-Neto, 2009 | Callichthys callichthys |
| Characiformes | Characidae | Achirus lineatus | native | marine | Andrade-Neto, 2009 | Achirus lineatus |
| Characiformes | Characidae | Acinocheirodon melanogramma | native | freshwater | Andrade-Neto, 2009 | Acinocheirodon melanogramma |
| Characiformes | Characidae | Aphyocheirodon sp. | native | freshwater | Report CEMIG, 2007 | - |
| Characiformes | Characidae | Astyanax bimaculatus | native | freshwater | Andrade-Neto, 2009 | Astyanax bimaculatus |
| Characiformes | Characidae | Astyanax brevirhinus | native | freshwater | Andrade-Neto, 2009 | Astyanax brevirhinus |
| Characiformes | Characidae | Astyanax cf. jequitinhonhae | native | freshwater | Report CEMIG, 2007 | Astyanax fasciatus |
| Characiformes | Characidae | Astyanax fasciatus | native | freshwater | Andrade-Neto, 2009 | Astyanax fasciatus |
| Characiformes | Characidae | Astyanax lacustris | native | freshwater | Pugedo et al., 2016 | Astyanax lacustris |
| Characiformes | Characidae | Astyanax scabripinnis | native | freshwater | Andrade-Neto, 2009 | Astyanax scabripinnis |
| Characiformes | Characidae | Astyanax sp. | native | freshwater | Report CEMIG, 2007 | - |
| Characiformes | Characidae | Astyanax turmalinensis | native | freshwater | Andrade-Neto, 2009 | Astyanax turmalinensis |
| Characiformes | Characidae | Hyphessobrycon cf. luetkeni | native | freshwater | Intertechne, 2009 | Hyphessobrycon luetkenii |
| Characiformes | Characidae | Hyphessobrycon sp. | native | freshwater | Report CEMIG, 2007 | - |
| Characiformes | Characidae | Knodus moenkhausii | | freshwater | Pugedo et al., 2016 | Knodus moenkhausii |
| Characiformes | Characidae | Mimagoniates sylvicola | native | freshwater | Andrade-Neto, 2009 | Mimagoniates sylvicola |
| Characiformes | Characidae | Moenkhausia costae | Introduced | freshwater | Andrade-Neto, 2009 | Moenkhausia costae |
| Characiformes | Characidae | Moenkhausia intermedia | native | freshwater | Intertechne, 2009 | Moenkhausia intermedia |
| Characiformes | Characidae | Nematocharax venustus | endangered | freshwater | Andrade-Neto, 2009 | Nematocharax venustus |
| Characiformes | Characidae | Oligosarcus hepsetus | native | freshwater | Andrade-Neto, 2009 | Oligosarcus hepsetus |
| Characiformes | Characidae | Oligosarcus macrolepis | native | freshwater | Andrade-Neto, 2009 | Oligosarcus macrolepis |
| Characiformes | Characidae | Serrapinnus zanatae | native | freshwater | Jerep, Camelier & Zanata, 2016 | Serrapinnus zanatae |
| Characiformes | Crenuchidae | Characidium cf. fasciatum | native | freshwater | Intertechne, 2009 | Characidium fasciatum |
| Characiformes | Crenuchidae | Characidium spp. | not described yet | freshwater | Andrade-Neto, 2009 | - |
| Characiformes | Curimatidae | Cyphocharax cf gilbert | native | freshwater | Intertechne, 2009 | Cyphocharax gilbert |
| Characiformes | Curimatidae | Cyphocharax jagunco | native | freshwater | Dutra et al., 2016 | Cyphocharax jagunco |
| Characiformes | Curimatidae | Cyphocharax lundi | native | freshwater | Dutra et al., 2016 | Cyphocharax lundi |
| Characiformes | Curimatidae | Cyphocharax naegelii | native | freshwater | Report CEMIG, 2005 | Cyphocharax naegelii |
| Characiformes | Curimatidae | Steindachnerina elegans | native | freshwater | Andrade-Neto, 2009 | Steindachnerina elegans |

| Order | Family | Species | Status | Habitat | Reference | Current status (Catalog of Fishes) |
|---|---|---|---|---|---|---|
| Characiformes | Erythrinidae | Hoplias brasiliensis | native | freshwater | Andrade-Neto, 2009 | Hoplias brasiliensis |
| Characiformes | Erythrinidae | Hoplias malabaricus | native | freshwater | Andrade-Neto, 2009 | Hoplias malabaricus |
| Characiformes | Prochilodontidae | Prochilodus argenteus | Introduced | freshwater | Sales et al., 2017 | Prochilodus argenteus |
| Characiformes | Prochilodontidae | Prochilodus costatus | Introduced | freshwater | Godinho et al., 1999 | Prochilodus costatus |
| Characiformes | Prochilodontidae | Prochilodus hartii | native | freshwater | Andrade-Neto, 2009 | Prochilodus hartii |
| Characiformes | Prochilodontidae | Prochilodus lineatus | Introduced | freshwater | Sales et al., 2017 | Prochilodus lineatus |
| Characiformes | Serrasalmidae | Colossoma macropomum | | freshwater | Pugedo et al., 2016 | Colossoma macropomum |
| Characiformes | Serrasalmidae | Serrasalmus brandtii | Introduced | freshwater | Pugedo et al., 2016 | Serrasalmus brandtii |
| Characiformes | Serrasalmidae | Serrasalmus sp. | Introduced | freshwater | Godinho, 2008 | - |
| Cichliformes | Cichlidae | Astronotus ocellatus | Introduced | freshwater | Bizerril & Lima, 2005 | Astronotus ocellatus |
| Cichliformes | Cichlidae | Australoheros sp. | | freshwater | Pugedo et al., 2016 | - |
| Cichliformes | Cichlidae | Cichla kelberi | Introduced | | Pugedo et al., 2016 | Cichla kelberi |
| Cichliformes | Cichlidae | Cichla sp. | Introduced | freshwater | FADETEC, 2002 | - |
| Cichliformes | Cichlidae | Cichlasoma facetum | native | freshwater | Report CEMIG, 2007 | Valid as Australoheros facetus |
| Cichliformes | Cichlidae | Crenicichla sp. | native | freshwater | Intertechne, 2009 | - |
| Cichliformes | Cichlidae | Geophagus brasiliensis | native | Freshwater; brackish | Andrade-Neto, 2009 | Geophagus brasiliensis |
| Cichliformes | Cichlidae | Oreochromis niloticus | Introduced | freshwater | Bizerril & Lima, 2005 | Oreochromis niloticus |
| Cichliformes | Cichlidae | Tilapia sp. | Introduced | freshwater | Godinho et al., 2001 | - |
| Clupeiformes | Engraulidae | Anchoviella lepidentostole | native | Marine; freshwater; brackish | Andrade-Neto, 2009 | Anchoviella lepidentostole |
| Clupeiformes | Engraulidae | Lycengraulis grossidens | native | Marine; freshwater; brackish | Andrade-Neto, 2009 | Lycengraulis grossidens |
| Cypriniformes | Cyprinidae | Hypophthalmichthys molitrix | Introduced | freshwater | Alves et al., 2007 | Hypophthalmichthys molitrix |
| Cyprinodontiformes | Poeciliidae | Phalloceros caudimaculatus | native | freshwater | Andrade-Neto, 2009 | Phalloceros caudimaculatus |
| Cyprinodontiformes | Poeciliidae | Phalloceros sp. | native | freshwater | Pugedo et al., 2016 | - |
| Cyprinodontiformes | Poeciliidae | Poecilia reticulata | Introduced | freshwater | Bizerril & Lima, 2005 | Poecilia reticulata |
| Cyprinodontiformes | Poeciliidae | Poecilia vivipara | native | freshwater | Intertechne, 2009 | Poecilia vivipara |
| Cyprinodontiformes | Rivulidae | Simpsonichthys espinhacensis | possibly endangered | freshwater | Nielsen, Pessali & Dutra, 2017 | Simpsonichthys espinhacensis |
| Cyprinodontiformes | Rivulidae | Simpsonichthys ocellatus | native | freshwater | Andrade-Neto, 2009 | Hypsolebias ocellatus |
| Cyprinodontiformes | Rivulidae | Simpsonichthys perpendicularis | endangered | freshwater | Andrade-Neto, 2009 | Ophthalmolebias perpendicularis |
| Gymnotiformes | Gymnotidae | Gymnotus bahianus | native | freshwater | Andrade-Neto, 2009 | Gymnotus bahianus |
| Gymnotiformes | Gymnotidae | Gymnotus carapo | native | freshwater | Andrade-Neto, 2009 | Gymnotus carapo |
| Gymnotiformes | Gymnotidae | Gymnotus pantherinus | native | freshwater | Andrade-Neto, 2009 | Gymnotus pantherinus |
| Gymnotiformes | Gymnotidae | Gymnotus sylvius | | | Pugedo et al., 2016 | Gymnotus sylvius |
| Gymnotiformes | Sternopygidae | Eigenmania virescens | native | freshwater | Andrade-Neto, 2009 | Eigenmania virescens |
| Mugiliformes | Mugilidae | Mugil platanus | native | Marine; freshwater; brackish | Andrade-Neto, 2009 | Synonym of Mugil liza |
| Perciformes | Carangidae | Caranx latus | native | Marine; freshwater; brackish | Andrade-Neto, 2009 | Caranx latus |
| Perciformes | Centropomidae | Centropomus parallelus | native | Marine; freshwater; brackish | Andrade-Neto, 2009 | Centropomus parallelus |
| Perciformes | Centropomidae | Centropomus undecimalis | native | Marine; freshwater; brackish | Andrade-Neto, 2009 | Centropomus undecimalis |
| Perciformes | Eleotridae | Dormitator maculatus | native | Marine; freshwater; brackish | Andrade-Neto, 2009 | Dormitator maculatus |
| Perciformes | Eleotridae | Eleotris pisonis | native | Marine; freshwater; brackish | Andrade-Neto, 2009 | Eleotris pisonis |

| Order | Family | Species | Status | Habitat | Reference | Current status (Catalog of Fishes) |
|---|---|---|---|---|---|---|
| Perciformes | Gerreidae | Eugerres brasilianus | native | marine | Pugedo et al., 2016 | Eugerres brasilianus |
| Perciformes | Gobiidae | Awaous tajasica | native | Freshwater; brackish; | Andrade-Neto, 2009 | Awaous tajasica |
| Siluriformes | Ariidae | Genidens genidens | native | marine;brackish | Andrade-Neto, 2009 | Genidens genidens |
| Siluriformes | Auchenipteridae | Pseudauchenipterus jequitinhonhae | native | freshwater | Andrade-Neto, 2009 | Pseudauchenipterus jequitinhonhae |
| Siluriformes | Auchenipteridae | Trachelyopterus galeatus | | freshwater | Pugedo et al., 2016 | Trachelyopterus galeatus |
| Siluriformes | Auchenipteridae | Trachelyopterus striatulus | native | freshwater | Andrade-Neto, 2009 | Trachelyopterus striatulus |
| Siluriformes | Callichthyidae | Aspidoras cf. rochai | native | freshwater | Report CEMIG, 2007 | Aspidoras cf. rochai |
| Siluriformes | Callichthyidae | Corydoras sp. | | freshwater | Pugedo et al., 2016 | - |
| Siluriformes | Callichthyidae | Hoplosternum littorale | | freshwater | Pugedo et al., 2016 | Hoplosternum littorale |
| Siluriformes | Clariidae | Clarias gariepinus | Introduced | freshwater | Godinho, 2007 | Clarias gariepinus |
| Siluriformes | Doradidae | Wertheimeria maculata | native | freshwater | Andrade-Neto, 2009 | Wertheimeria maculata |
| Siluriformes | Heptapteridae | Imparfinis sp. | native | freshwater | Intertechne, 2009 | - |
| Siluriformes | Heptapteridae | Pariolius sp. | native | freshwater | Report CEMIG, 2007 | - |
| Siluriformes | Heptapteridae | Pimelodella sp. | not described yet | freshwater | Andrade-Neto, 2009 | - |
| Siluriformes | Heptapteridae | Rhamdia jequitinhonha | endangered | freshwater | Andrade-Neto, 2009 | Rhamdia jequitinhonha |
| Siluriformes | Heptapteridae | Rhamdia quelen | native | freshwater | Andrade-Neto, 2009 | Rhamdia quelen |
| Siluriformes | Loricariidae | Chauliocheilos saxatilis | native | freshwater | Martins et al., 2014 | Chauliocheilos saxatilis |
| Siluriformes | Loricariidae | Delturus brevis | native | freshwater | Andrade-Neto, 2009 | Delturus brevis |
| Siluriformes | Loricariidae | Harttia garavelloi | native | freshwater | Andrade-Neto, 2009 | Harttia garavelloi |
| Siluriformes | Loricariidae | Hypostomus nigrolineatus | native | freshwater | Zawadzki et al., 2016 | Hypostomus nigrolineatus |
| Siluriformes | Loricariidae | Hypostomus sp. | not described yet | freshwater | Andrade-Neto, 2009 | - |
| Siluriformes | Loricariidae | Microlepidogaster discus | native | freshwater | Martins et al., 2014 | Microlepidogaster discus |
| Siluriformes | Loricariidae | Pareiorhaphis sp. | native | freshwater | Report CEMIG, 2007 | - |
| Siluriformes | Loricariidae | Pareiorhaphis stephanus | native | freshwater | Andrade-Neto, 2009 | Pareiorhaphis stephana |
| Siluriformes | Loricariidae | Pareiorhaphis lineata | native | freshwater | Pereira et al., 2017 | Pareiorhaphis lineata |
| Siluriformes | Loricariidae | Parotocinclus jequi | native | freshwater | Lehman et al., 2013 | Parotocinclus jequi |
| Siluriformes | Loricariidae | Parotocinclus sp. | native | freshwater | Report CEMIG, 2007 | - |
| Siluriformes | Loricariidae | Pogonopoma wertheimeri | native | freshwater | Andrade-Neto, 2009 | Pogonopoma wertheimeri |
| Siluriformes | Pimelodidae | Pseudoplatystoma spp. | Introduced | freshwater | Godinho, 2007 | - |
| Siluriformes | Pimelodidae | Steindachneridion amblyurum | endangered | freshwater | Andrade-Neto, 2009 | Steindachneridion amblyurum |
| Siluriformes | Trichomycteridae | Trichomycterus itacambirussu | native | freshwater | Andrade-Neto, 2009 | Trichomycterus itacambirussu |
| Siluriformes | Trichomycteridae | Trichomycterus jequitinhonhae | native | freshwater | Andrade-Neto, 2009 | Trichomycterus jequitinhonhae |
| Siluriformes | Trichomycteridae | Trichomycterus landinga | native | freshwater | Andrade-Neto, 2009 | Trichomycterus landinga |
| Siluriformes | Trichomycteridae | Trichomycterus spp. | not described yet | freshwater | Report CEMIG, 2007 | - |
| Siluriformes | Pseudopimelodidae | Microglanis cf. parahybae | native | freshwater | Intertechne, 2009 | Microglanis parahybae |
| Synbranchiformes | Synbranchidae | Synbranchus marmoratus | native | freshwater;brackish | Andrade-Neto, 2009 | Synbranchus marmoratus |

Table S2: List of samples sequenced for the custom reference database.

| Sample | Species |
|---|---|
| 233 | *Astyanax bimaculatus* |
| 242 | *Astyanax fasciatus* |
| 226 | *Astyanax fasciatus* |
| 6584 | Australoeros sp. |
| 6585 | Australoeros sp. |
| 1897 | *Brycon aff. devillei* |
| 4508 | *Brycon ferox* |
| 3745 | *Brycon opalinus* |
| 3679 | *Brycon opalinus* |
| 3678 | *Brycon opalinus* |
| 3062 | Brycon sp. |
| 3063 | Brycon sp. |
| 4635 | Brycon sp. 2 |
| 4636 | Brycon sp. 2 |
| 66 | Brycon sp. |
| 1619 | Characidium sp. |
| NAI103 | Characidium sp. |
| 3683 | *Characidium timbuiense* |
| 294 | *Cichlasoma facetum* |
| 293 | *Cichlasoma facetum* |
| 1573 | Corydoras sp. |
| 3556 | *Crenicichla lacustris* |
| NAI4579 | *Crenicichla lacustris* |
| NAI3555 | *Crenicichla lacustris* |
| 3549 | *Cyphocharax gilbert* |
| 1810 | *Cyphocharax gilbert* |
| 3549 | *Cyphocharax gilbert* |
| 1830 | *Delturus carinotus* |
| 1832 | *Delturus carinotus* |
| 304 | *Geophagus brasiliensis* |
| 302 | *Geophagus brasiliensis* |
| 303 | *Geophagus brasiliensis* |
| 440 | *Harttia garavelloi* |
| 441 | *Harttia garavelloi* |
| 3703 | Hasemania sp. |
| 1164 | Hisonotus sp. |
| 104 | *Hoplias brasiliensis* |
| 3573 | *Hoplias intermedius* |
| 3714 | *Hoplias intermedius* |
| 258 | *Hoplias malabaricus* |
| 259 | *Hoplias malabaricus* |
| 2644 | *Hoplosternum litoralle* |
| 2643 | *Hoplosternum litoralle* |
| 1873 | *Hypomasticus garmani* |
| NAI3713 | *Hypomasticus mormyrops* |
| 1566_B | *Hypoptopomatinae* |
| 1565 | *Hypoptopomatinae* |
| 1566 | *Hypoptopomatinae* |
| 4211 | *Hypostomus gr. affinis* |
| 4212 | *Hypostomus gr. affinis* |
| 3712 | Hypostomus sp. |
| 1069 | Hypostomus sp. |
| 3544 | *Leporinus copelandii* |
| 1823 | *Leporinus copelandii* |
| NAI1823 | *Leporinus copelandii* |
| NAI3033 | *Leporinus copelandii* |
| 296 | *Leporinus crassilabris* |
| 221 | *Leporinus crassilabris* |
| 84 | *Leporinus elongatus* |
| 85 | *Leporinus elongatus* |
| 52 | *Leporinus garmani* |
| NAI107 | *Leporinus garmani* |
| 58 | *Leporinus garmani* |
| 93 | *Leporinus steindachneri* |
| 2672 | *Loricariichthys castaneus* |
| BB run | *Loricariichthys castaneus* |
| 236 | *Moenkhausia costae* |
| NAI305 | *Moenkhausia costae* |
| 1822 | Neoplecostominae |
| 4519 | *Oligosarcus argenteus* |
| NAI3547 | *Oligosarcus argenteus* |
| NAI1818 | *Oligosarcus argenteus* |
| 4555 | *Oligosarcus macrolepis* |
| 1621 | Pareiorhaphis sp. |
| 3710 | Pareiorhaphis sp. |
| NAI | Pareiorhaphis sp. |
| 467 | *Pareiorhaphis stephanus* |
| 466 | *Pareiorhaphis stephanus* |
| 4144 | *Phalloceros elachistos* |
| NAI4144 | *Phalloceros elachistos* |
| NAI3704 | Phalloceros sp. |
| 3704 | Phalloceros sp. |
| 1629 | Pimelodella sp. |
| NAI116 | *Prochilodus harttii* |
| 3056B | Rhamdia cf. jequitinhonha |
| 3056A | Rhamdia cf. jequitinhonha |
| 3055 | Rhamdia cf. jequitinhonha |
| 3056 | Rhamdia cf. jequitinhonha |
| NAI101 | *Serrasalmus brandtii* |
| 2996 | *Steindachneridion amblyurum* |
| 2996 | *Steindachneridion amblyurum* |
| 3014 | *Steindachneridion amblyurum* |
| 232 | *Steindachnerina elegans* |
| 6556 | *Trachelyopterus striatulus* |
| 6583 | *Trachelyopterus striatulus* |
| 1833 | *Trachelyopterus striatulus* |
| NAI1833 | *Trachelyopterus striatulus* |
| 1620 | Trichomycterus sp.1 |
| 1624 | Trichomycterus sp.2 |
| NAI3708 | Trichomycterus sp. |
| 5778 | Trichomycterus sp. |
| 77 | *Wertheimeria maculata* |
| 76 | *Wertheimeria maculata* |

Table S3: Comparison of different SWARM clustering thresholds.

| | Sampling event 1 | | | Sampling event 2 | | |
|---|---|---|---|---|---|---|
| | LIB1 | | | LIB2 | | |
| | **d=1** | **d=2** | **d=3** | **d=1** | **d=2** | **d=3** |
| **Number of swarms** | 107567 | 60259 | 41094 | 131121 | 70633 | 46339 |
| **Largest swarm** | 64783 | 135807 | 179901 | 40359 | 52581 | 91237 |
| **Max generations** | 28 | 30 | 31 | 21 | 24 | 29 |
| | **After SWARM Recount** | | | | | |
| **Clusters** | 107567 | 60259 | 41094 | 131121 | 70633 | 46339 |
| **Cluster >5 reads** | 5249 | 4563 | 3595 | 5714 | 4766 | 3823 |
| **Reads kept for calculations/Total reads = 404914** | 302596 | 349218 | 367415 | 369295 | 428835 | 452186 |
| **Alignment cached** | 81.44% | 78.15% | 75.67 | 79.34 | 74.66 | 70.02 |
| **Number of MOTUs** | 5249 | 4563 | 3595 | 5714 | 4766 | 3823 |
| **Number of reads** | 3955997 | 4004093 | 4022641 | 3914479 | 3975772 | 3999426 |
| **Actinopterygii-MOTUs** | 4821 | 3974 | 3065 | 5054 | 4128 | 3265 |
| **Neotropical Orders-MOTUs** | 3195 | 2534 | 2168 | 2421 | 2041 | 1691 |
| **Neotropical Families-MOTUs** | 2635 | 2058 | 1862 | 1732 | 1441 | 1007 |
| **Neotropical Species-MOTUs** | 2056 | 1664 | 1491 | 967 | 893 | 820 |
| **Number of MOTUs assigned to species (minid 0.97)** | 155 | 52 | 25 | 59 | 42 | 28 |
| **Number of MOTUs assigned to genus (minid 0.97)** | 58 | 16 | 5 | 64 | 20 | 6 |
| **Number of MOTUs assigned to Family (minid 0.97)** | 22 | 7 | 0 | 25 | 5 | 2 |
| **Number of MOTUs assigned to Suborder (minid 0.97)** | 4 | 2 | 1 | 2 | 0 | 0 |

Table S4: Species identified applying the minimum identity of 0.97, according to each SWARM threshold.

| MOTUs assigned to species (minid 0.97) | | | | | | | | | | | |
|---|---|---|---|---|---|---|---|---|---|---|---|
| First Sampling event | | | | | | Second Sampling event | | | | | |
| LIB1 | | | | | | LIB2 | | | | | |
| d=1 | | d=2 | | d=3 | | d=1 | | d=2 | | d=3 | |
| Species | Number of MOTUs | Species | Number of MOTUs | Species | Number of MOTUs | Species | Number of MOTUs | Species | Number of MOTUs | Species | Number of MOTUs |
| Astronotus ocellatus | 1 | Astronotus ocellatus | 1 | Astronotus ocellatus | 1 | Astronotus ocellatus | 1 | Astronotus ocellatus | 1 | Astronotus ocellatus | 1 |
| Australoheros facetus | 1 | Australoheros facetus | 1 | Australoheros facetus | 1 | Brycon sp. | 1 | Brycon sp. | 1 | Characidium sp. | 1 |
| Brycon sp. | 1 | Crenicichla lacustris | 6 | Crenicichla lacustris | 3 | Characidium sp. | 6 | Coptodon zillii | 1 | Coptodon zillii | 1 |
| Characidium sp. | 11 | Cyphocharax gilbert | 1 | Cyprinus carpio | 1 | Coptodon zillii | 1 | Crenicichla lacustris | 2 | Crenicichla lacustris | 2 |
| Crenicichla lacustris | 9 | Cyprinus carpio | 1 | Delturus carinotus | 1 | Crenicichla lacustris | 3 | Cyphocharax gilbert | 1 | Cyphocharax gilbert | 1 |
| Cyphocharax gilbert | 2 | Delturus carinotus | 2 | Geophagus brasiliensis | 1 | Cyphocharax gilbert | 1 | Delturus carinotus | 1 | Delturus carinotus | 1 |
| Cyprinus carpio | 1 | Geophagus brasiliensis | 1 | Gymnotus carapo | 1 | Delturus carinotus | 1 | Geophagus brasiliensis | 1 | Geophagus brasiliensis | 1 |
| Delturus carinotus | 6 | Gymnotus carapo | 1 | Hoplias malabaricus | 1 | Geophagus brasiliensis | 1 | Gymnotus carapo | 1 | Gymnotus carapo | 1 |
| Geophagus brasiliensis | 6 | Hoplias malabaricus | 1 | Hoplosternum littorale | 1 | Gymnotus carapo | 1 | Hoplias intermedius | 1 | Hoplias intermedius | 1 |
| Gymnotus carapo | 1 | Hoplosternum littorale | 1 | Hypomasticus mormyrops | 1 | Hoplias intermedius | 1 | Hoplias malabaricus | 1 | Hoplias malabaricus | 1 |
| Hoplias malabaricus | 1 | Hypomasticus mormyrops | 4 | Lophiosilurus alexandri | 1 | Hoplias malabaricus | 1 | Hoplosternum littorale | 1 | Hoplosternum littorale | 1 |
| Hoplosternum littorale | 6 | Hypostomus gymnorhynchus | 1 | | 2 | Hoplosternum littorale | 1 | Hypomasticus mormyrops | 1 | Hypomasticus mormyrops | 1 |
| Hypomasticus mormyrops | 6 | Hypostomus nigromaculatus | 1 | Oreochromis aureus | 1 | Hypomasticus mormyrops | 1 | Hypostomus nigromaculatus | 1 | Lophiosilurus alexandri | 1 |
| Hypostomus gymnorhynchus | 1 | Leporinus copelandii | 3 | Phalloceros sp.J | 1 | Hypostomus nigromaculatus | 1 | Leporinus copelandii | 2 | Megaleporinus garmani | 1 |
| Hypostomus nigromaculatus | 1 | Lophiosilurus alexandri | 1 | Poecilia reticulata | 1 | Leporinus copelandii | 3 | Lophiosilurus alexandri | 1 | Moenkhausia costae | 1 |
| Leporinus copelandii | 14 | Megaleporinus garmani | 1 | Salminus brasiliensis | 1 | Lophiosilurus alexandri | 2 | Megaleporinus garmani | 2 | Neoplecostominae gen. 2 sp. FFR-2012 | 1 |
| Lophiosilurus alexandri | 5 | Moenkhausia costae | 3 | Trachelyopterus striatulus | 1 | Megaleporinus garmani | 9 | Moenkhausia costae | 3 | Neoplecostomini gen.n. sp.n TEP-2017 | 1 |
| Megaleporinus garmani | 14 | Neoplecostominae gen. 2 sp. FFR-2012 | 1 | Trichomycterus sp. | 1 | Moenkhausia costae | 3 | Neoplecostominae gen. 2 sp. FFR-2012 | 2 | Oligosarcus argenteus | 1 |
| Moenkhausia costae | 6 | Neoplecostomini gen.n. sp.n TEP-2017 | 3 | Trichomycterus sp.J | 1 | Neoplecostominae gen. 2 sp. FFR-2012 | 2 | Neoplecostomini gen.n. sp.n TEP-2017 | 1 | Oreochromis aureus | 1 |
| Neoplecostominae gen. 2 sp. FFR-2012 | 3 | Oreochromis aureus | 1 | Wertheimeria maculata | 1 | Neoplecostomini gen.n. sp.n TEP-2017 | 1 | Oligosarcus argenteus | 1 | Poecilia reticulata | 1 |
| Neoplecostomini gen.n. sp.n TEP-2017 | 6 | Phalloceros sp.J | 1 | | | Oligosarcus argenteus | 1 | Oreochromis aureus | 1 | Rhamdia quelen | 1 |
| Oligosarcus argenteus | 3 | Poecilia reticulata | 1 | | | Oreochromis aureus | 1 | Phalloceros sp.J | 1 | Salminus brasiliensis | 1 |
| Oreochromis aureus | 1 | Prochilodus argenteus | 2 | | | Phalloceros sp.J | 1 | Poecilia reticulata | 1 | Serrasalmus brandtii | 1 |
| Phalloceros sp.J | 2 | Rhamdia quelen | 1 | | | Poecilia reticulata | 1 | Prochilodus argenteus | 4 | Trachelyopterus striatulus | 1 |
| Poecilia reticulata | 1 | Salminus brasiliensis | 2 | | | Prochilodus argenteus | 3 | Rhamdia quelen | 2 | Trichomycterus sp.J | 1 |
| Prochilodus argenteus | 28 | Trachelyopterus striatulus | 1 | | | Rhamdia quelen | 2 | Salminus brasiliensis | 2 | Wertheimeria maculata | 1 |
| Rhamdia quelen | 1 | Trichomycterus sp. | 1 | | | Salminus brasiliensis | 1 | Serrasalmus brandtii | 2 | | |
| Salminus brasiliensis | 5 | Trichomycterus sp.J | 2 | | | Serrasalmus brandtii | 3 | Trachelyopterus striatulus | 1 | | |
| Serrasalmus brandtii | 1 | Wertheimeria maculata | 1 | | | Trachelyopterus striatulus | 1 | Trichomycterus sp.J | 1 | | |
| Trachelyopterus striatulus | 5 | | | | | Trichomycterus sp.J | 3 | Wertheimeria maculata | 1 | | |
| Trichomycterus sp. | 1 | | | | | Wertheimeria maculata | 1 | | | | |
| Trichomycterus sp.J | 1 | | | | | | | | | | |
| Wertheimeria maculata | 1 | | | | | | | | | | |

Table S5: Taxa detected by eDNA metabarcoding and correspondent nearest neighbour species reported for Jequitinhonha river basin.

| Order | Family | Genus | eDNA taxon | Nearest neighbor species reported for JRB | | | |
|---|---|---|---|---|---|---|---|
| Characiformes | Anostomidae | Hypomasticus | *Hypomasticus mormyrops* | *Megaleporinus garmani)* | | | |
| | | Leporinus | *Leporinus copelandii* | *Leporinus bahiensis* | *Leporinus* spp. | *Leporinus steindachneri* | *Leporinus taeniatus* |
| | | Megaleporinus | *Megaleporinus garmani* | *Megaleporinus garmani* | *Megaleporinus elongatus* | | |
| | Bryconidae | Brycon | *Brycon* sp. | *Brycon devillei* | *Brycon* sp. | | |
| | | Salminus | *Salminus brasiliensis** | | | | |
| | Characidae | Moenkhausia | *Moenkhausia costae* | *Moenkhausia costae* | *Moenkhausia intermedia* | | |
| | | Oligosarcus | *Oligosarcus argenteus* | *Oligosarcus hepsetus* | *Oligosarcus macrolepis* | | |
| | Crenuchidae | Characidium | *Characidium* sp. | *Characidium spp.* | *Characidium* cf. *fasciatum* | | |
| | Curimatidae | Cyphocharax | *Cyphocharax gilbert* | *Cyphocharax jagunco* | *Cyphocharax lundi* | *Cyphocharax naegelii* | |
| | Erythrinidae | Hoplias | *Hoplias malabaricus* | *Hoplias brasiliensis* | *Hoplias malabaricus* | | |
| | Prochilodontidae | Prochilodus | *Prochilodus argenteus* | *Prochilodus hartii* | *Prochilodus argenteus* | *Prochilodus costatus* | *Prochilodus lineatus* |
| | Serrasalmidae | Serrasalmus | *Serrasalmus brandtii* | *Serrasalmus brandtii* | *Serrasalmus* sp. | | |
| Cichliformes | Cichlidae | Astronotus | *Astronotus ocellatus* | *Astronotus ocellatus* | | | |
| | | Australoheros | *Australoheros facetus* | *Australoeros* sp. | | | |
| | | Coptodon | *Coptodon zillii* | Tilapia sp. | | | |
| | | Crenicichla | *Crenicichla lacustris* | *Crenicichla* sp. | | | |
| | | Geophagus | *Geophagus brasiliensis* | *Geophagus brasiliensis* | | | |
| | | Oreochromis | *Oreochromis aureus* | *Oreochromis niloticus* | | | |
| Cypriniformes | Cyprinidae | Cyprinus | *Cyprinus carpio* | *Hypophthalmichthys molitrix* | | | |
| Cyprinodontiformes | Poeciliidae | Phalloceros | *Phallocero* s sp. | *Phalloceros caudimaculatus* | *Phalloceros* sp. | | |
| | | Poecilia | *Poecilia reticulata* | *Poecilia reticulata* | *Poecilia vivipara* | | |
| Gymnotiformes | Gymnotidae | Gymnotus | *Gymnotus carapo* | *Gymnotus bahianus* | *Gymnotus carapo* | *Gymnotus pantherinus* | *Gymnotus sylvius* |
| Siluriformes | Auchenipteridae | Trachelyopterus | *Trachelyopterus striatulus* | *Trachelyopterus galeatus* | *Trachelyopterus striatulus* | | |
| | Callichthyidae | Hoplosternum | *Hoplosternum littorale* | *Hoplosternum littorale* | | | |
| | Doradidae | Wertheimeria | *Wertheimeria maculata* | *Wertheimeria maculata* | | | |
| | Heptapteridae | Rhamdia | *Rhamdia quelen* | *Rhamdia quelen* | *Rhamdia jequitinhonha* | | |
| | Loricariidae | Delturus | *Delturus carinotus* | *Delturus brevis* | | | |
| | | Hypostomus | *Hypostomus gymnorhynchus* | *Hypostomus nigrolineatus* | *Hypostomus* sp. | | |
| | | | *Hypostomus nigromaculatus* | *Hypostomus nigrolineatus* | *Hypostomus* sp. | | |
| | | | Neoplecostominae gen. 2 sp. FFR-2012 | | | | |
| | | | Neoplecostomini gen.n. sp.n TEP-2017 | | | | |
| | Pseudopimelodidae | Lophiosilurus | *Lophiosilurus alexandri** | | | | |
| | Trichomycteridae | Trichomycterus | *Trichomycterus* sp. | *Trichomycterus itacambirussu* | *Trichomycterus jequitinhonhae* | *Trichomycterus landinga* | *Trichomycterus* spp. |
| | | | *Trichomycterus* sp.2 | *Trichomycterus itacambirussu* | *Trichomycterus jequitinhonhae* | *Trichomycterus landinga* | *Trichomycterus* spp. |

* Species not previously reported for JRB

Table S6: Taxa detected in each sampling event and sampling medium.

| **WATER** | | **SEDIMENT** | |
|---|---|---|---|
| **Sampling event 1** | **Sampling event 2** | **Sampling event 1** | **Sampling event 2** |
| *Astronotus ocellatus* | *Astronotus ocellatus* | *Characidium* sp. | *Astronotus ocellatus* |
| *Australoheros facetus* | *Brycon* sp. | *Crenicichla lacustris* | *Brycon* sp. |
| *Brycon* sp. | *Characidium* sp. | *Cyphocharax gilbert* | *Characidium* sp. |
| *Characidium* sp. | *Coptodon zillii* | *Delturus carinotus* | *Coptodon zillii* |
| *Crenicichla lacustris* | *Crenicichla lacustris* | *Geophagus brasiliensis* | *Crenicichla lacustris* |
| *Cyphocharax gilbert* | *Cyphocharax gilbert* | *Hoplias malabaricus* | *Cyphocharax gilbert* |
| *Cyprinus carpio* | *Delturus carinotus* | *Hoplosternum littorale* | *Delturus carinotus* |
| *Delturus carinotus* | *Geophagus brasiliensis* | *Hypomasticus mormyrops* | *Geophagus brasiliensis* |
| *Geophagus brasiliensis* | *Gymnotus carapo* | *Hypostomus nigromaculatus* | *Gymnotus carapo* |
| *Gymnotus carapo* | *Hoplias intermedius* | *Leporinus copelandii* | *Hoplias intermedius* |
| *Hoplias malabaricus* | *Hoplias malabaricus* | *Lophiosilurus alexandri* | *Hoplosternum littorale* |
| *Hoplosternum littorale* | *Hoplosternum littorale* | *Megaleporinus garmani* | *Hypomasticus mormyrops* |
| *Hypomasticus mormyrops* | *Hypomasticus mormyrops* | *Moenkhausia costae* | *Hypostomus nigromaculatus* |
| *Hypostomus gymnorhynchus* | *Hypostomus nigromaculatus* | *Neoplecostominae* gen. 2 sp. FFR-2012 | *Leporinus copelandii* |
| *Hypostomus nigromaculatus* | *Leporinus copelandii* | *Neoplecostomini* gen.n. sp.n TEP-2017 | *Lophiosilurus alexandri* |
| *Leporinus copelandii* | *Lophiosilurus alexandri* | *Oligosarcus argenteus* | *Megaleporinus garmani* |
| *Lophiosilurus alexandri* | *Megaleporinus garmani* | *Oreochromis aureus* | *Moenkhausia costae* |
| *Megaleporinus garmani* | *Moenkhausia costae* | *Phalloceros* sp. | *Neoplecostominae* gen. 2 sp. FFR-2012 |
| *Moenkhausia costae* | *Neoplecostominae* gen. 2 sp. FFR-2012 | *Prochilodus argenteus* | *Neoplecostomini* gen.n. sp.n TEP-2017 |
| *Neoplecostominae* gen. 2 sp. FFR-2012 | *Neoplecostomini* gen.n. sp.n TEP-2017 | *Rhamdia quelen* | *Oligosarcus argenteus* |
| *Neoplecostomini* gen.n. sp.n TEP-2017 | *Oligosarcus argenteus* | *Salminus brasiliensis* | *Oreochromis aureus* |
| *Oligosarcus argenteus* | *Oreochromis aureus* | *Serrasalmus brandtii* | *Phalloceros* sp. |
| *Oreochromis aureus* | *Phalloceros sp.* | *Trachelyopterus striatulus* | *Prochilodus argenteus* |
| *Phalloceros* sp. | *Poecilia reticulata* | *Trichomycterus* sp. | *Rhamdia quelen* |
| *Poecilia reticulata* | *Prochilodus argenteus* | *Wertheimeria maculata* | *Salminus brasiliensis* |
| *Prochilodus argenteus* | *Rhamdia quelen* | | *Serrasalmus brandtii* |
| *Rhamdia quelen* | *Salminus brasiliensis* | | *Trachelyopterus striatulus* |
| *Salminus brasiliensis* | *Serrasalmus brandtii* | | *Trichomycterus* sp. |
| *Serrasalmus brandtii* | *Trachelyopterus striatulus* | | *Wertheimeria maculata* |
| *Trachelyopterus striatulus* | *Trichomycterus* sp. | | |
| *Trichomycterus* sp. | *Wertheimeria maculata* | | |
| *Trichomycterus* sp. | | | |
| *Wertheimeria maculata* | | | |

Table S7: Sample sites including code, city, human population* and GPS coordinates.

| Site | City | Population | Coordinates | |
|---|---|---|---|---|
| 1 | Mendanha | 639 | 18° 7'15.06"S | 43°30'59.16"W |
| 2 | Terra Branca | <1000 | 17°18'48.34"S | 43°12'26.61"W |
| 3 | Jose Gonçalves | 4553 | 16°52'37.8"S | 42°43'51.9"W |
| 4 | Itacambiruçu | 15024 | 16°36'24.00"S | 42°49'46.00"W |
| 5 | Coronel Murta | 9117 | 16°44'26.85"S | 42°34'11.78"W |
| 6 | Araçuaí | 36013 | 16°52'27.9"S | 42°06'59.3"W |
| 7 | Jequitinhonha | 24131 | 16°25'35.8"S | 41°00'52.8"W |
| 8 | Almenara | 38755 | 16° 8'26.20"S | 40°35'4.64"W |
| 9 | Salto da Divisa | 6859 | 15°59'51.07"S | 39°53'29.76"W |
| 10 | Itapebi | 10495 | 15°56'57.69"S | 39°31'27.08"W |
| 11 | Belmonte | 21798 | 15°51'0.02"S | 38°52'13.66"W |

*Human population census based on IBGE, 2018
